# Primary multistep phosphorelay activation comprises both cytokinin and abiotic stress responses in Brassicaceae

**DOI:** 10.1101/2023.11.14.567013

**Authors:** Katrina Leslie Nicolas Mala, Jan Skalak, Elena Zemlyanskaya, Vladislav Dolgikh, Veronika Jedlickova, Helene S. Robert-Boisivon, Lenka Havlicková, Klara Panzarova, Martin Trtilek, Ian Bancroft, Jan Hejatko

## Abstract

Multistep phosphorelay (MSP) signaling integrates hormonal and environmental signals to control plant development and adaptive responses. The type-A *RESPONSE REGULATORs* (*RRAs*), the downstream members of the MSP cascade and cytokinin primary response genes, are supposed to mediate primarily the negative feedback regulation of (cytokinin-induced) MSP signaling. However, the transcriptional data suggest the involvement of *RRAs* in stress-related responses as well. By employing evolutionary conservation with the well-characterized *Arabidopsis thaliana RRAs*, we identified 5 and 38 novel putative *RRAs* in *Brassica oleracea* and *Brassica napus*, respectively. Our phylogenetic analysis suggests the existence of gene-specific selective pressure, maintaining the homologs of *ARR3, ARR6,* and *ARR16* as singletons during the evolution of *Brassica oleracea* and *Brassica rapa*. We categorized *RRAs* based on the kinetics of their cytokinin-mediated upregulation and observed both similarities and specificities in this type of response across Brassicaceae. Using bioinformatic analysis and experimental data demonstrating the cytokinin responsiveness of *Arabidopsis*-derived *TCSv2* reporter, we unveil the mechanistic conservation of cytokinin-mediated upregulation of *RRAs* in *Brassica rapa* and *Brassica napus*. Notably, we identify partial cytokinin dependency of cold stress-induced *RRA* transcription, thus corroborating the role of cytokinin signaling in the crop adaptive responses.

**Highlights:** We identified *Brassica* homologs of *Arabidopsis* type-A response regulators (*RRAs*), demonstrate existence of selective pressure preventing several *RRAs* multiplication during Brassicaceae evolution and describe cytokinin dependency of cold-induced *RRAs* upregulation.

## Introduction

Cytokinins regulate a wide range of biological processes that are vital for plant growth and development (Cortleven *et al*., 2019; Werner and Schmulling, 2009; Zurcher and Muller, 2016). In *Arabidopsis thaliana*, cytokinin signaling occurs through a multistep phosphorelay (MSP), sometimes also called two-component signaling (Kieber and Schaller, 2018). The core components of MSP include ARABIDOPSIS HISTIDINE KINASEs (AHKs), ARABIDOPSIS HISTIDINE-CONTAINING PHOSPHOTRANSMITTERs (AHPs), and ARABIDOPSIS RESPONSE REGULATORs (RRs). In the presence of cytokinins, the CHASE-containing AHKs (AHK2, AHK3, and AHK4) located at the plasma membrane or endoplasmic reticulum (ER) undergo autophosphorylation at a conserved His residue and transfer the phosphate group to the conserved Asp residue within the AHK receiver domain (Antoniadi *et al*., 2020; Hwang and Sheen, 2001; Inoue *et al*., 2001; Kubiasova *et al*., 2020; Muller and Sheen, 2007). Cytoplasmic AHPs accept the phosphate from the AHKs and translocate to the nucleus, allowing the final transphosphorylation of the receiver domain of type-B RRs (RRBs) and transcriptional regulation of the cytokinin-responsive genes.

Besides the aforementioned RRBs, the *A. thaliana* genome contains two more types of RRs: type-A RRs (RRAs) and type-C RRs [RRCs; (Imamura *et al*., 1998; Schaller *et al*., 2008)]. RRBs possess a cytokinin-responsive receiver domain along with a large C-terminal extension that harbors the GARP (Golden/ARR/Psr1) motif, a Myb-like DNA binding domain (Hosoda *et al*., 2002). In contrast, the RRAs are characterized by the presence of a receiver domain and short C-terminal sequences but do not contain the DNA-binding domain. *RRAs* act as cytokinin primary response genes, being rapidly induced by cytokinins via direct transcriptional activation by RRBs, even in the absence of *de novo* protein synthesis (D’Agostino *et al*., 2000; Taniguchi *et al*., 1998). RRA proteins are phosphorylated by RRBs and mediate the negative regulation of MSP signaling via yet-unknown mechanisms (Lee *et al*., 2008). There are ten known *RRAs* in *A. thaliana* (*ARR3, ARR4, ARR5, ARR6, ARR7, ARR8, ARR9, ARR15, ARR16,* and *ARR17*), acting as partially redundant negative regulators of (cytokinin-induced) MSP signaling (To *et al*., 2004). Previous studies have demonstrated the key role of *A. thaliana RRAs* in several developmental and growth regulatory processes including stem cell specification, meristem activity, and regeneration (Buechel *et al*., 2010; Leibfried *et al*., 2005; Muller and Sheen, 2008; Zhao *et al*., 2010).

The transcriptional activity of *RRAs* was shown to be linked to diverse abiotic stress responses, including salinity, cold, and drought (Bhaskar *et al*., 2021; Jain *et al*., 2006; Jeon *et al*., 2010; Kang *et al*., 2012; Sharan *et al*., 2017; Shi *et al*., 2012; Tran *et al*., 2007; Urao *et al*., 1998; Wang *et al*., 2019). However, compared to *A. thaliana*, the role of *RRAs* in the cytokinin and stress responses remains rudimentary for a majority of agronomically important plant species. Advancements in molecular biology, particularly sequencing technologies, have facilitated the genome-wide identification of putative components of the MSP cascade not only in *A. thaliana* (Hwang and Sheen, 2001) but also in crop species such as rice (Ito and Kurata, 2006; Jain *et al*., 2006; Karan *et al*., 2009; Pareek *et al*., 2006; Sharan *et al*., 2017; Tsai *et al*., 2012), maize (Asakura *et al*., 2003), soybean (Mochida *et al*., 2010), and wheat (Sun *et al*., 2022). Genes involved in MSP signaling have been reported in Chinese cabbage (*B. rapa* spp. Pekinensis) (Kaltenegger *et al*., 2018; Liu *et al*., 2014), *B. oleracea* (Kaltenegger *et al*., 2018) and *B. napus* (Jiang et al., 2022; Kuderova et al., 2015). Diploid *B. oleracea, B. rapa,* and allotetraploid *B. napus* species are among the most commercially valuable crops in the Brassicaceae family. They are not only consumed as culinary vegetables but also cultivated as oilseed crops, covering approximately 38 million hectares in 2021/2022 in various countries, including Canada, China, India, European Union, and Australia (European Commision, 2019; Kumar *et al*., 2009; Rathore *et al*., 2022).

In this study, we identify novel *RRAs* in *B. napus* and *B. oleracea* and provide insights into the evolutionary relationships, kinetics, and mechanism of cytokinin responses, as well as the involvement of cytokinin in the abiotic stress-mediated modulation of *RRAs* within the *A. thaliana* and *Brassica* species.

## Materials and methods

### Identification of type-A response regulators in *Brassica* species, motif search, multiple sequence alignment, and chromosomal mapping

The protein sequences of the 10 known type A RRs in the *Arabidopsis thaliana* genome (Hwang *et al*., 2002) were obtained from NCBI (https://www.ncbi.nlm.nih.gov/protein/) (NCBI reference sequence ARR3 NP_176202.1, ARR4 NP_001321924.1, ARR5 NP_190393.1, ARR6 NP_201097.1, ARR7 NP_173339.1, ARR8 NP_181663.1, ARR9 NP_001325622.1, ARR15 NP_177627.1, ARR16 NP_181599.1, ARR17 NP_567037.1) (Supplementary Table S1). These sequences were used as queries in Protein BLAST (BLASTP) searches against the protein database of *B. oleracea, B. rapa,* and *B. napus* in the EnsemblPlants (Release 51) (Howe *et al*., 2021). Genes were selected as described by Kaltenegger *et al*. (2018). The coding sequences, genomic sequences, and protein sequences of the selected genes were retrieved from EnsemblPlants (Release 51) (Howe *et al*., 2021) and Brassicaceae Database (BRAD version 3.0; http://brassicadb.cn) (Chen *et al*., 2021).

Using the Expassy SIM-Alignment Tool for protein sequences with BLOSUM62 as a comparison matrix (https://web.expasy.org/sim/) (Duvaud *et al*., 2021), the amino acid sequence homology of the identified *Brassica* RRAs was compared with *A. thaliana* RRAs (Supplementary Table S2). Similarly, the *B. napus RRAs* from both A and C subgenomes were compared to their progenitor species *B. rapa* and *B. oleracea.* The presence of the conserved response regulator domain was analyzed using the GenomeNet Bioinformatics Tools, sequence motif search, MOTIF (https://www.genome.jp/tools/motif/) of Kyoto University Bioinformatics Center (Kyoto University Bioinformatics Center, 2015). The protein sequences of the identified Brassica *RRAs* were used as input, and a search against the PFAM database was performed with a cut-off score of E-value of 1. Sequences that possessed the conserved response regulator receiver domain (Rec) (PF00072) were selected for further analysis in this study.

Multiple sequence alignment was conducted using the MUSCLE algorithm (Edgar, 2004) implemented in the UGENE (Okonechnikov *et al*., 2012a) to annotate the location of important conserved residues. The genomic locations of *A. thaliana* and *Brassica RRAs* were retrieved from the EnsemblPlants (Release 51) (Howe *et al*., 2021) and BrassicaDB (BRAD version 3.0; http://brassicadb.cn) databases (Chen *et al*., 2021). These locations were visualized using MapGene2Chrom (MG2C_v2.1, http://mg2c.iask.in/mg2c_v2.1/) (Chao *et al*., 2015) by setting appropriate parameters for the figure output. The identified Brassica *RRA* genes were named following the nomenclature proposed by Heyl *et al*. (2013), and the numbers assigned to them correspond to their *A. thaliana* counterparts after performing the phylogenetic analysis. In cases where multiple homologs of *ARR4*, *ARR5*, *ARR7*, *ARR8*, *ARR9*, *ARR15*, and *ARR17* were found in Brassica, they were designated with the letters “a”, “b”, or “c” following descending order of homology depending on the percentage of amino acid identities they share with that specific *RRA*.

### Phylogenetic analysis of type A response regulator genes and gene structure analysis

A comparative phylogenetic analysis was conducted using MEGA7: Molecular Evolutionary Genetics Analysis version 7.0 for bigger datasets (Kumar *et al*., 2016) based on the alignment of the conserved response regulator domain (Rec) (PF00072) as described by Kaltenegger *et al*. (2018). The multiple sequence alignment was performed using the conserved Rec domain using the MUSCLE algorithm (Edgar, 2004) implemented in MEGA7 (Kumar *et al*., 2016). The Neighbor-Joining method (Saitou and Nei, 1987) was used to infer the evolutionary history. The evolutionary distances were computed using the Poisson correction method (Zuckerkandl and Pauling, 1965) and are expressed as the number of amino acid substitutions per site. The analysis included 1000 bootstrap replicates, and all ambiguous positions were removed for each sequence pair. Phylogenetic trees were constructed to compare the individual *Brassica* species with *A. thaliana* RRAs, as well as to compare all the Brassica RRAs among themselves. Gene structure analysis of *A. thaliana* and *Brassica* RRAs including their schematic representations was made using Gene Structure Display Server (http://gsds.gao-lab.org/) (Hu *et al*., 2015).

Dual synteny plots were created using the TBTools dual synteny plot function (Chen *et al*., 2020) to compare the *Brassica* species with *A. thaliana*, and *B. napus* with its parental species, *B. rapa* and *B. oleracea.* Before plotting the dual synteny, a one-step MCScanX analysis was performed in TBTools. The genome sequence files and gene structure annotation files for *Brassica* species and *A. thaliana* were retrieved from EnsemblPlants (Release 54) (Cunningham *et al*., 2021).

### Plant materials, hormones, and abiotic stress treatment

Seeds of *A. thaliana* (Col-0), *B. rapa* (R-0-18), *B. oleracea* (DH1012), and *B. napus* (Darmor) were cultivated on 1/2 MS media for one week inside the growth chamber under controlled conditions. Before cultivation, the seeds underwent a cold pre-treatment in darkness at 4°C for 3 days. The growth chamber was maintained at a temperature of 21^◦^C /18^◦^C for a 16-hour day/8-hour night photoperiod, with 130 µE light intensity.

To investigate the expression profile of the 10 *A. thaliana RRAs* and 66 *Brassica RRAs* after cytokinin treatment, one-week-old seedlings were exposed to exogenous treatment with 5 µM BAP for 0 hour, 0.5 hour, 1 hour, 2 hours, and 4 hours as described (D’Agostino *et al*., 2000).

For the abiotic stress treatment, cold-treated seedlings were incubated at 4°C in the presence of white light. Seedlings subjected to salinity stress were treated with 250 mM NaCl, while osmotic stress was induced using a 300 mM mannitol solution. All stress treatments were applied for 0 hour, 2 hours, and 4 hours.

Additionally, a separate cold treatment experiment was conducted following the methodology described above to assess the expression of cold-responsive *ARR7*, and its *Brassica* homologs. The focus of this experiment was to evaluate the effects of PI-55, a known antagonist of the cytokinin receptor activity (Spichal *et al*., 2009). One-week-old seedlings were treated with either PI-55 (0.1 µM/1 µM) or DMSO and incubated either under cold (4°C) or control conditions (21^◦^C) for 4 hours.

### RNA Isolation and RT-qPCR Analysis

Total RNA was extracted from the collected seedlings following the Quick-Start Protocol included in the RNeasy® Plant Mini Kit (QIAGEN, Germany). Additionally, DNAse treatment was performed using an RNase-Free DNase set (QIAGEN) to remove any DNA contamination. The concentration, integrity, and purity of the extracted RNA samples were examined using NanoDrop One UV spectrophotometer (Thermo Fisher Scientific). Reverse transcription was performed to generate first-strand cDNA using the SuperScript^TM^ III First-Strand Synthesis System (ThermoFisher Scientific) with 1 µg of RNA using oligo-dT primer. For the expression profiling of *RRAs* after cytokinin treatment and abiotic stress exposure, 66 out of the 78 *Brassica RRAs* along with the 10 *A. thaliana RRAs* were analyzed. For the expression profiling of cold-responsive *ARRs* after PI-55 treatment, *ARR7* and its *Brassica* homologs (i.e., *BrRRA7b, BoRRA7a, BoRRA7b, BnARRA7a, BnARRA7b, BnCRRA7a,* and *BnCRRA7b)* were analyzed. Several reference genes were utilized as an internal control, including the commonly used housekeeping genes (Guénin *et al*., 2009) such as *UBQ10* and *UBC10* (added for abiotic stress) for *Arabidopsis*, *BrELF1* for *B. rapa*, *BoTUB6* for *B. oleracea*, *BnACT2A* and *BnACT2C* for *B. napus* (primers listed in Supplementary Table S3). All primers used were designed based on the following features: product size (70–200bp), primer length (18–22bp), Tm (59–65°C), GC content (50–60%), target gene specificity, and absence of nucleotide repeats. The reactions for RT-qPCR were performed using FastStart SYBR® Green Master (Roche Diagnostics GmbH) on the Rotor-Gene Q 5plex HRM Platform (QIAGEN, Germany). Melting curve analysis was performed to confirm the specificity of the product for each primer pair. The relative gene expression level was calculated relative to the control using the delta-delta Ct method (Pfaffl, 2004). The RT-qPCR analysis was performed in three independent biological replicates, each with three technical replicates. Subsequently, a heatmap representation of the expression of type A*RRAs* after exogenous cytokinin treatment and abiotic stress treatment was generated and presented as the log2 fold-change (log_2_FC). The heatmap was constructed using Cluster 3.0 for Windows (de Hoon *et al*., 2004) and viewed using Java TreeView (Saldanha, 2004).

### Analysis of *cis*-regulatory elements in the promoter regions of type A *RR* genes across *Brassica* species

Multiple sequence alignment of the homologous RRB amino acid sequences from *Brassica* species and *A. thaliana* was performed using Clustal Omega (Madeira *et al*., 2022) to assess the conservation of their GARP-like DNA binding domains. The alignment was visualized using the MView online tool (Madeira *et al*., 2022). Reference genomes and genome annotations for *A. thaliana*, *B. rapa*, *B. oleracea*, and *B. napus* were downloaded from EnsemblPlants (Yates *et al*., 2022). The upstream regulatory sequences of protein-coding genes were extracted from the reference genomes using GFF3 annotations with the Bedtools getfasta tool (Quinlan and Hall, 2010). The publicly available ChIP-seq data for *A. thaliana* transcription factors (TFs) ARR1 and ARR10 (Xie *et al*., 2018) was used for a *de novo* motif search with Homer (Heinz *et al*., 2010). To identify potential RRB binding sites in gene regulatory regions, the Position Weight Matrices (PWMs) were used. The thresholds for PWMs were calculated using the previously described algorithm (Touzet and Varré, 2007). Then the PWMs were applied to three 500-bp-long intervals of protein-coding genes: [−1500; −1000], [−1000; −500], [−500; +1] relative TSS. To compare the density of potential RRB binding sites in the regulatory regions of *Brassica RRA* coding genes (used in the cytokinin and abiotic stress treatment) to random expectation (which is the density of the binding sites in the regulatory regions of all protein-coding genes), Fisher’s exact test was used. To account for multiple testing, we used Bonferroni correction: *p*-value threshold was set as 0.05/24. The fold enrichment was calculated as the ratio of RRB binding site density in RRA regulatory regions to the average density in the corresponding regions of all protein-coding genes.

The promoter sequences of *A. thaliana* and *Brassica RRAs* (used in the cytokinin and abiotic stress treatment) were also subjected to *insilico* analysis using the online database, PlantCARE (http://bioinformatics.psb.ugent.be/webtools/plantcare/html/) (Lescot *et al*., 2002). The objective was to investigate the presence of environmental stress-responsive *cis*-elements in these sequences. Additionally, a Pearson correlation analysis was conducted to determine the relationship between the gene expression of cold-responsive *A. thaliana RRAs* (*ARR6*, *ARR7*, and *ARR15*) and Brassica *RRAs* (*BrRRA6, BrRRA7a, BrRRA7b, BrRRA15a, BrRRA15b, BoRRA6, BoRRA7a, BoRRA7b, BoRRA15a, BoRRA15b, BnARRA6, BnARRA7a, BnARRA7b, BnARRA15a, BnCRRA6, BnCRRA7a, BnCRRA7b, BnCRRA15a, BnCRRA15b*) after 2 hours and 4 hours of cold exposure, and the total number of environmental stress-related *cis*-elements within the promoter regions of these genes. In the case of *A. thaliana*, additional comparisons were made using the DAPseq data to select transcription factors (TFs) with potential binding sites in the *A. thaliana* promoters. Moreover, to assess the enrichment of the TF binding sites, particularly the position weight matrix (PWM) models in *A. thaliana*, a comparison was made between stress-sensitive promoters and stress-insensitive promoters for both *A. thaliana* and *Brassica* species.

### Transformation of *Brassica* species with *TCSv2*:3XVENUS and CK treatment

The *TCSv2*:3XVENUS construct, obtained from Maya Barr (Steiner *et al*., 2020), was subcloned into the pGREEN0029 binary vector (Hellens *et al*., 2000) and introduced into *B. rapa* (R-0-18), *B. oleracea* (DH1012), and *B. napus* (Darmor), following the protocol described by Jedlickova *et al*. (2022). Only root tips of *B. rapa* and *B. napus* transformed hairy roots were used in the experiment, as the transformation for *B. oleracea* was unsuccessful. Root tips of *B. rapa* and *B. napus* hairy roots were gathered 2 weeks after subculturing and treated with either 5 µM synthetic 6-benzylaminopurine (BAP) or 0.1 % DMSO for 0 hour, 0.5 hour, 1 hour, 2 hours, and 4 hours, as described (D’Agostino *et al*. (2000).

### Root imaging, and quantification of reporter gene expression

Root tips were imaged using the laser scanning confocal imaging microscope Zeiss LSM780 Axio-Observer, equipped with an external In Tune laser (488-649 nm, <3 nm width, pulsed at 40 MHz, 1.5 mW C-Apochromat) and a 20x objective. The expression of VENUS in the root apical meristem (RAM) was quantified using IMAGEJ software (Schneider *et al*., 2012) and the spot detection algorithm in IMARIS 9.0 (Bitplane, http://www.bitplane.com/imaris/imaris). Representative images generated using IMARIS are presented. To ensure accurate analysis, the fluorescence intensity of each DMSO or BAP-treated root was initially normalized to the area of the scanned roots (in pixels) and further normalized to the fluorescence intensity of the roots at the start of the treatment (0 h). Subsequently, the relative fluorescence intensity was calculated as the ratio of normalized fluorescence intensity in BAP-treated roots to the normalized fluorescence intensity of DMSO-treated roots.

### Statistical analysis

A one-way ANOVA followed by Dunnett’s test was conducted to evaluate differences in the calculated relative fluorescence intensity in the scanned roots at the start and after 0.5 hour, 1 hour, 2 hours, and 4 hours of exogenous BAP treatment. Furthermore, a two-way ANOVA followed by Tukey’s HSD multiple comparison test was employed to compare the relative expression of cold-responsive *ARR7, BrRRA7a, BrRRA7b, BoRRA7a, BoRRA7b, BnARRA7a, BnARRA7b, BnCRRA7a,* and *BnCRRA7b* after PI-55 treatment. All statistical analysis was conducted using the GraphPad Prism version 9.0 for Windows (GraphPad Software, San Diego, California USA, www.graphpad.com).

## Results

### The type-A response regulators and their genomic distribution in the Brassicaceae family

Using a similarity search (see Materials and Methods for more details), we identified 78 putative *RRAs* in *B. oleracea*, *B. rapa,* and *B. napus* that share a high degree of sequence identity with *A. thaliana RRAs* (Fig. 1 and Supplementary Table S2). Among these, 20 were previously reported by Kaltenegger *et al*. (2018) and Liu *et al*. (2014) in the genome of *B. rapa* and following previously agreed nomenclature (Heyl *et al*., 2013), we designated them as *BrRRAs* (Fig. 1B). In the genome of *B. oleracea,* we found 20 putative *RRAs* (designated as *BoRRAs*), including 5 novel putative *RRAs* that were not included in Kaltenegger *et al*. (2018) (Fig. 1C). Lastly, we recognized 38 novel putative *RRAs* in *B. napus*, 20 of which located in the A subgenome (*BnARRAs*) and 18 in the C subgenome (*BnCRRAs*) (Fig. 1D). The putative paralogues were indexed with “a”, “b” or “c” in an order following the decreasing percentage of amino acid identities they share with the corresponding *RRAs* from *A. thaliana*.

**Figure 1.**
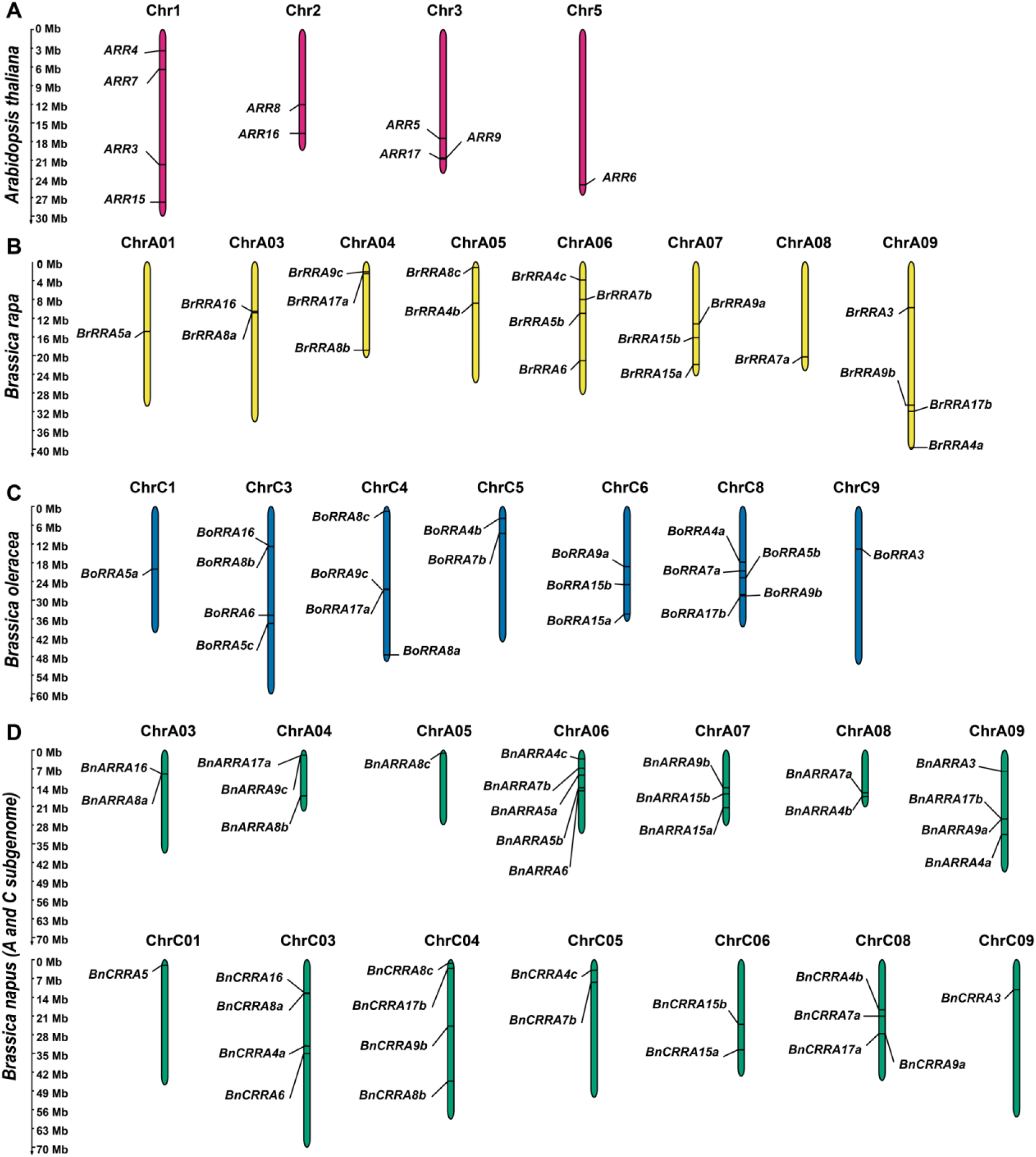
Chromosomal localization of known and newly identified *RRAs* in *Brassicaceae*. *RRAs* in **(A)** *Arabidopsis thaliana*, **(B)** *Brassica rapa*, **(C)** *Brassica oleracea*, and **(D)** *Brassica napus*. The figures were generated using MapGene2Chrom (MG2C_v2.1, http://mg2c.iask.in/mg2c_v2.1/) (Chao *et al*., 2015).

*BrRRAs* were mapped to chromosomes ChrA01, ChrA03, ChrA04, ChrA05, ChrA06, ChrA07, ChrA08, and ChrA09, while *BoRRAs* located on ChrC1, ChrC3, ChrC4, ChrC5, ChrC6, ChrC8, and ChrC9 (Fig. 1B, C). As expected, *BnARRAs* and *BnCRRAs* were found to localize to corresponding homologous chromosomes in A and C subgenomes, respectively (ChrA03, ChrA04, ChrA05, ChrA06, ChrA07, ChrA08, ChrA09 for *BnARRAs* and ChrC01, ChrC03, ChrC04, ChrC05, ChrC06, ChrC08, ChrC09 for *BnCRRAs*; Fig. 1D).

### *Brassica* and *A. thaliana* RRAs show a high level of conservation

A motif search in the putative protein sequences of all the 78 *Brassica* RRAs confirmed the presence of the conserved Rec domain harboring the highly conserved D-<u>D</u>-K motif, including the (underlined) phosphoaccepting Asp, which is essential for the role of RRAs in mediating the negative feedback regulation of cytokinin signaling (Lee *et al*., 2008) (Fig. 2A, B). Moreover, all the predicted *Brassica* RRAs had protein sizes comparable with their putative *A. thaliana* orthologues (identified based on their phylogenetic analysis, see later in the text and Fig. 3), ranging from 127 to 265 amino acid residues, with ARR4 and ARR17 and their homologs being the longest and shortest, respectively (Fig. 2 and Supplementary Table S2). The evolutionary relationship among the RRAs (78 *Brassica* and 10 *A. thaliana* RRAs) was assayed by aligning the amino acid sequences of the conserved Rec domains (Fig. 3A, Supplementary Fig. S1 - S3). As expected, we observed a high level of conservation between the RRAs from *Brassica* and *A. thaliana*. The tree consists of five main clades, each composed of two subclades, reflecting the presence of five couples of very similar/paralogous RRAs (ARR7/ARR15, ARR5/ARR6, ARR3/ARR4, ARR16/ARR17 and ARR8/ARR9). This information was used to designate the individual *Brassica* RRAs according to their clustering into individual paralogous subclades (Fig. 3A).

**Figure 2.**
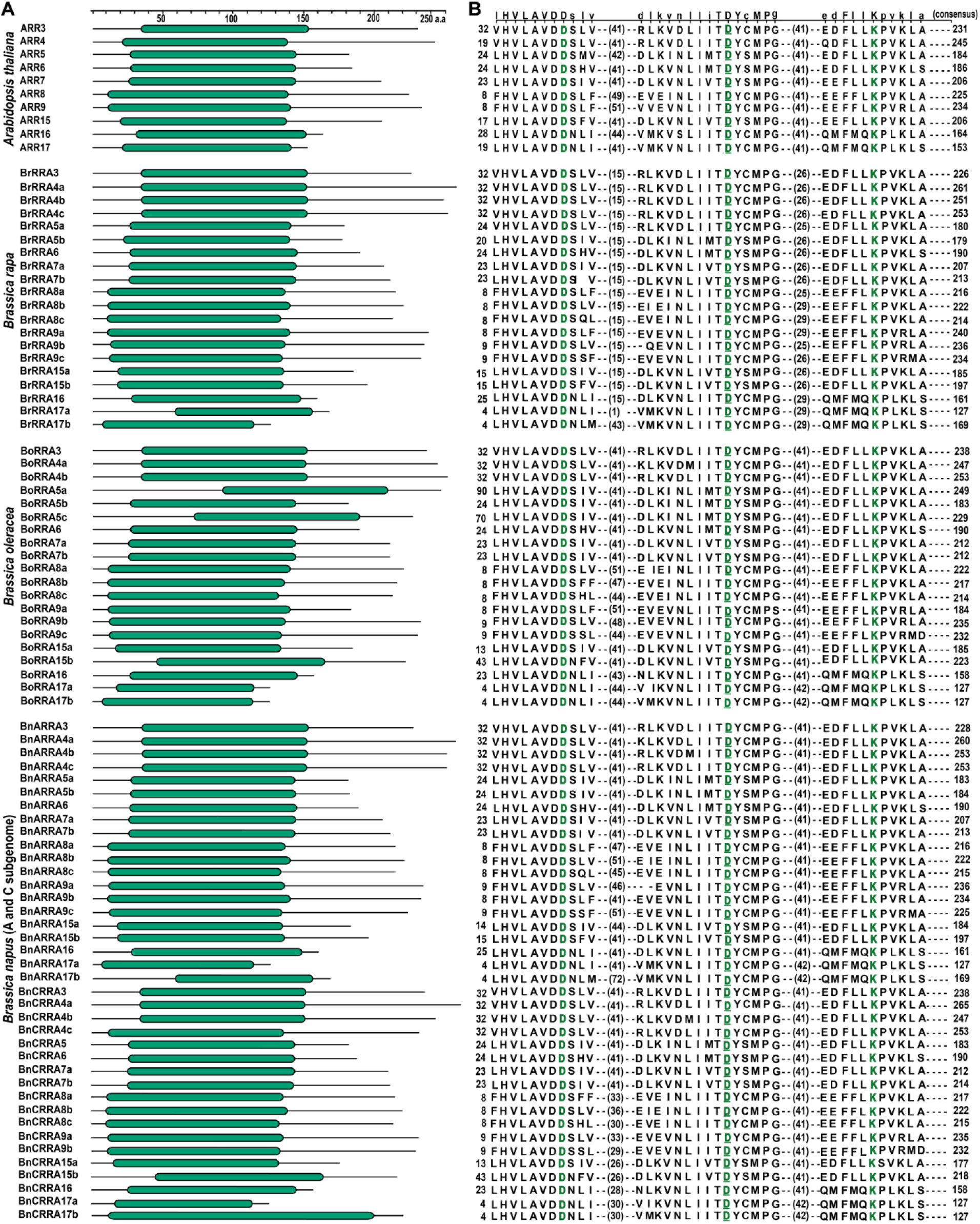
*A. thaliana* and *Brassica RRAs* reveal a high level of domain structure and amino acid sequence conservation. **(A)** Schematic depiction of the receiver domain (Rec, green rounded rectangle) localization in *A. thaliana* and *Brassica* RRAs identified using MOTIF (https://www.genome.jp/tools/motif/). **(B)** Amino acid sequence consensus within the conserved Rec domain (top line) based on the multiple alignments of the individual RRA protein sequences by MUSCLE algorithm in UGENE (Edgar, 2004; Okonechnikov *et al*., 2012b); the conserved D-D-K motif is highlighted in green.

**Figure 3.**
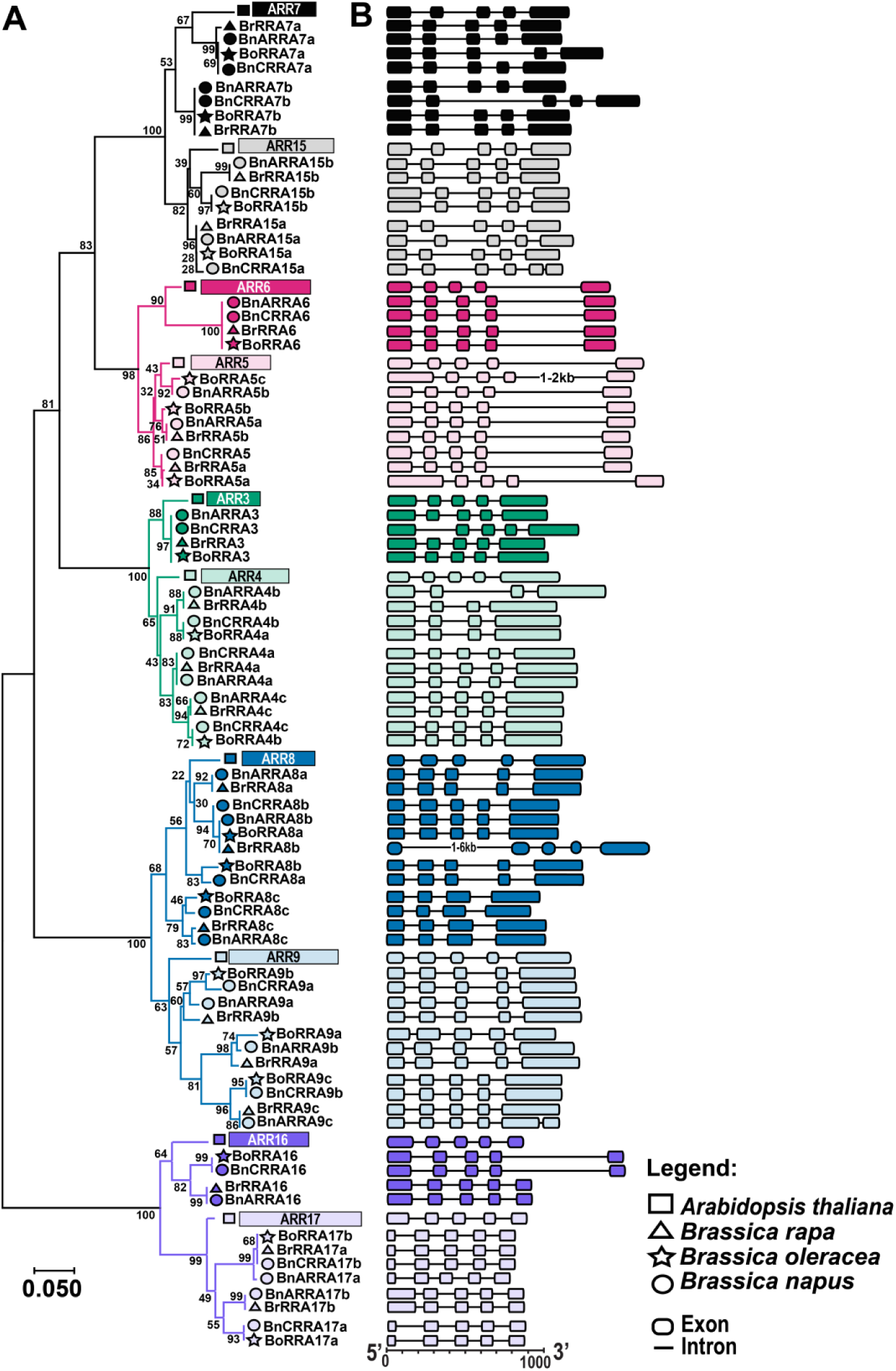
Phylogenetic relationships and gene structures of *RRAs* in Brassicaceae. **(A)** The unrooted tree is based on the similarity of RRA Rec domains constructed using the neighbor-joining method in MEGA 7 (Kumar *et al*., 2016); the bar represents the relative divergence of the examined sequences. The subclades composed of RRAs potentially orthologous to individual *A. thaliana RRAs* were colored using the same color; the subclades comprising homologs of the paired *A. thaliana RRAs*, the result of α WGD event (see the main text for details) are distinguished by different shades of a given color. The RRAs from individual species are distinguished by triangle (BrRRAs), star (BoRRAS), and circle (BnRRAs). **(B)** A schematic representation of the *A. thaliana* and *Brassica RRA* gene structures (exons are depicted as boxes separated by introns as lines) constructed using the Gene Structure Display Server (GSDS2.0 http://gsds.gao-lab.org/index.php) (Hu *et al*., 2015); the color code used as in (A).

The analysis of gene structure revealed that, except for 8 *RRAs* containing only 4 exons (*BrRRA4b, BoRRA4a, BnARRA4b, BnCRRA4b, BoRRA8c, BrRRA8c, BnCRRA8c,* and *BnARRA8c*), all other *RRAs* shared a gene model consisting of 5 exons and 4 introns (Fig. 3B). Among these, *ARR6* and its *Brassica* homologs exhibited nearly identical gene structures, including the number and length of exons and introns. Furthermore, genome-to-genome synteny analysis between the individual *Brassica* species and *A. thaliana* revealed that 20 out of 20 (20/20) *BrRRAs*, 11 out of 20 *BoRRAs*, and 32 out of 38 *BnRRAs* genes were syntenic with their *A. thaliana* counterparts (Fig. 4A). In case of *B. napus*, 36 out of 38 *BnRRAs* were syntenic with those of *B. rapa* and *B.* oleracea (Fig. 4B).

**Figure 4.**
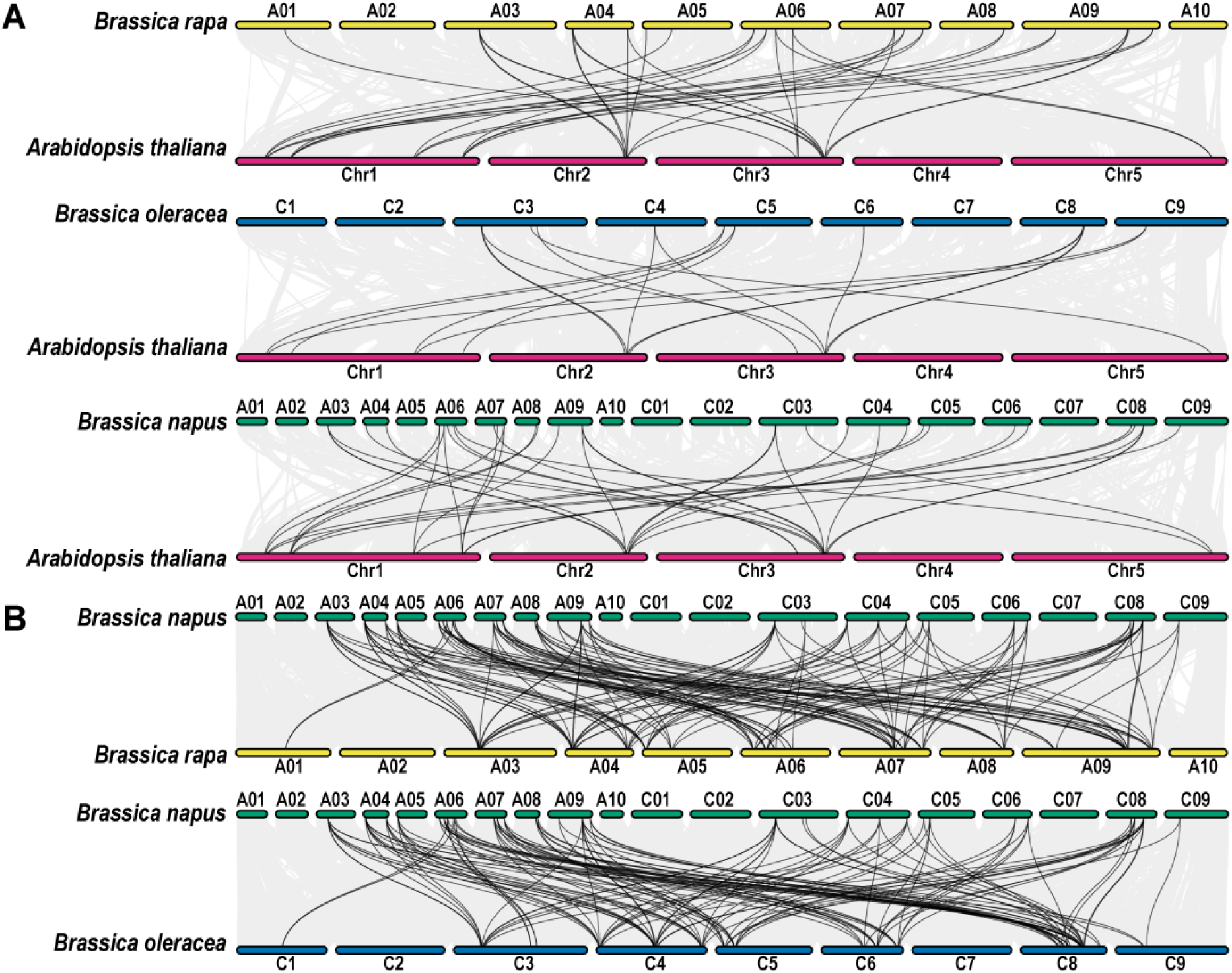
The synteny analysis of *Brassica RRAs*. **(A)** The synteny plots of the *RRAs* from assayed *Brassica* species with *A. thaliana*. **(B)** The synteny plots between *Brassica napus* and its parental species, *B. rapa*, and *B. oleracea*. Gray lines represent the syntenic blocks between the species, while the black lines represent the syntenic *RRA* pairs within the compared genomes.

Taken together, a high level of amino acid sequence conservation was observed within the *Brassica* species, confirming the previously described evolutionary relationships (Hendriks *et al*., 2022; Cheng *et al*., 2012; Cheng *et al*., 2014; Morinaga, 1929; Nagaharu and Nagaharu, 1935; Nikolov *et al*., 2019).

### Cytokinin treatment revealed the shared and distinct patterns of the *RRA* expression profiles between *A. thaliana* and *Brassica*

The *A. thaliana RRAs* are considered primary cytokinin response genes, as their transcription is promptly induced by exogenous cytokinins even in the absence of *de novo* protein synthesis (D’Agostino *et al*., 2000; Taniguchi *et al*., 1998). To compare the effects of cytokinin on the expression of *A. thaliana* and *Brassica RRAs*, one-week-old *A. thaliana* and *Brassica* seedlings were exposed to exogenous cytokinins for various time points ranging from 30 min to 4 hours (Fig. 5 and Supplementary Table S4).

**Figure 5.**
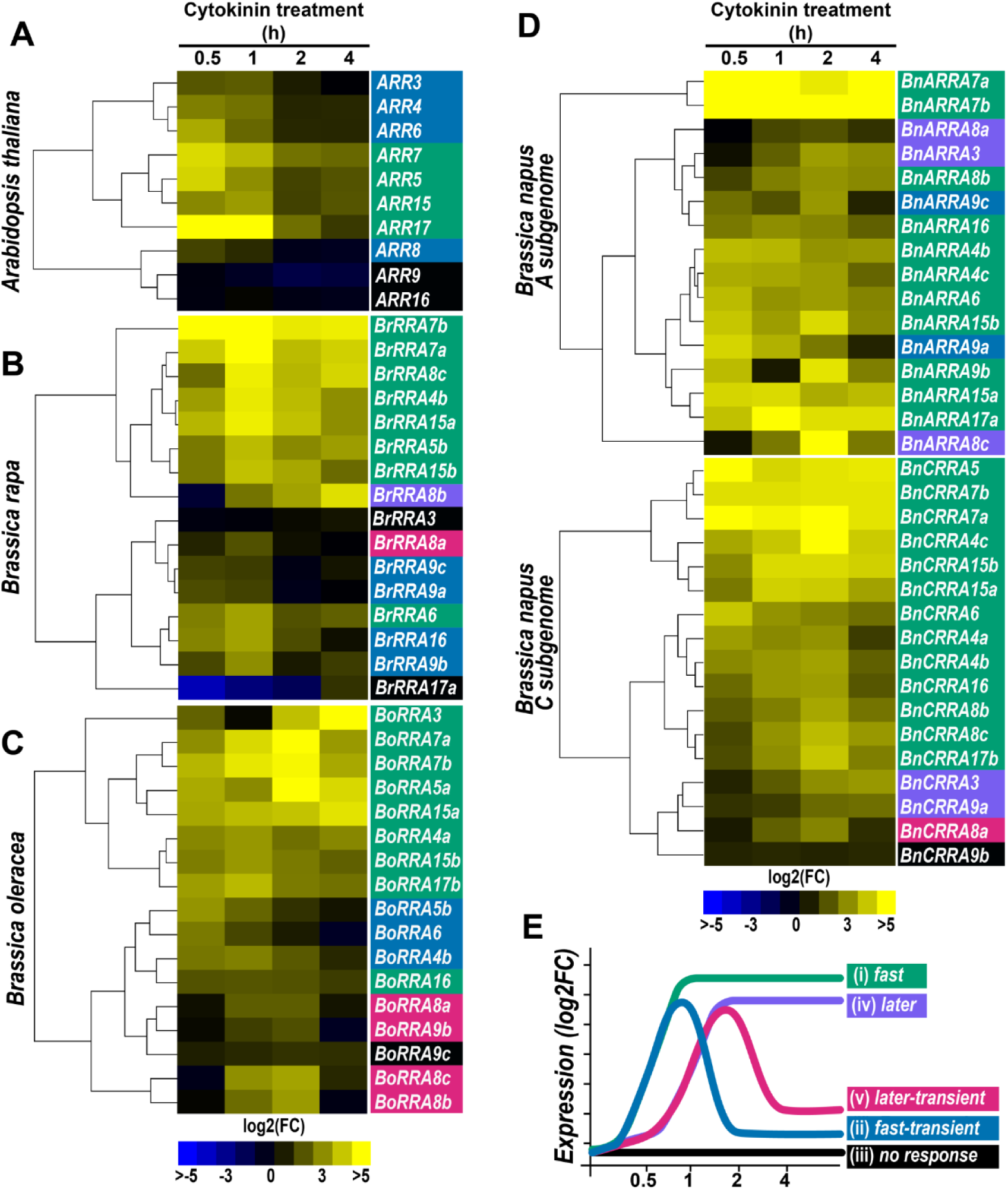
Kinetics of *A. thaliana* and *Brassica RRAs* response to cytokinins. Heatmaps representing the relative change of *RRA* expression in the one-week-old seedlings after cytokinin (5 µM BAP) treatment for the given time (0.5, 1, 2, and 4 h) normalized to mock-treated controls in **(A)** *A. thaliana*, **(B)** *Brassica rapa*, **(C)** *B. oleracea,* and **(D)** *B. napus*. The expression data is presented as log2 of fold-change between BAP- and mock-treated samples normalized by the delta-delta Ct (Pfaffl, 2004). **(E)** Schematic depiction of identified expression profile categories. The categorization of individual RRAs in panel A-D is color-coded as defined in (E).

Based on the timecourse of the observed transcriptional response, the expression profiles of individual *A. thaliana RRAs* were classified into three categories: *(i) fast,* exhibiting prompt upregulation after 30 min of cytokinin treatment followed by a gradual decline of expression throughout the rest of the treatment period, *(ii) fast-transient,* similar to *(i)*, but revealing fast decline after the initial peak and *(iii) no response* (Fig. 5A, E and Supplementary Table S4). In *A. thaliana*, we observed the same number (four) of *RRAs* with cytokinin response profiles classified as *fast* and *fast-transient* and two *RRAs* belonging to the *no response* category (Fig. 5A). In contrast, in *B. rapa* and *B. oleracea*, the proportion of *RRAs* with the *fast* profile increased at the expense of the *fast-transient* and two additional categories emerged: *(iv) later*, characterized by delayed upregulation occurring after 1 hour of cytokinin treatment and persisting until 4 hours, and *(v) later-transient*, similar to the *later* category but with a decline in expression at 4 hours (Fig. 5B, C, E). The decrease in the number of *RRAs* of the *fast-transient* category was more pronounced in *B. oleracea* compared to *B. rapa.* This trend was even more evident when comparing the A and C subgenome-specific *RRAs* in *B. napus,* where at least two *RRAs* of the fast-transient profile were still retained among the *BnARRAs* (encoded by the A subgenome of *B. rapa* origin), but no *fast-transient RRA* profile was found among *BnCRRAs* (located in the C subgenome originating from *B. oleracea*; compare Fig. 5B, C, D). In the case of *BrRRA17a,* we observed quite strong downregulation early after cytokinin application; however, for the sake of simplicity, *BrRRA17a* was categorized as belonging to category *(iii) no response* (Fig. 5B).

Analyzing the cytokinin response of individual *RRA* across the *Brassica* species and *A. thaliana*, similar expression profiles were observed for *ARR5, ARR7,* and *ARR15,* and most of their homologs in *B. rapa*, *B. oleracea*, and *B. napus*. However, a higher level of expression change (log_2_FC) of these *RRAs* was observed in the *Brassica* species compared to *A. thaliana* and this trend was apparent, particularly for *B. napus* homologs of *ARR7* (Fig. 5 and Supplementary Table S4). That aligns with RNA-sequencing profiling results of *B. napus* cultivars using the Renewable Industrial Products from Rapeseed (RIPR) diversity panel (Havlickova *et al*., 2018), which identified *ARR7* orthologues as one of the most abundant *RRAs* among the *B. napus* cultivars (Supplementary Fig. S4).

To wrap it up, all assayed *RRAs* across the Brassicaceae family were upregulated by cytokinins, demonstrating partially overlapping, but also species-specific temporal expression patterns.

### RRBs mediate the cytokinin-induced upregulation of *Brassica RRAs*

In *Arabidopsis*, cytokinin-dependent transcriptional activation of *RRAs* is mediated by RRBs, the cytokinin-regulated transcription factors that bind specific *cis*-regulatory motif enriched in the promoters of cytokinin-responsive genes (Muller and Sheen, 2008). To assess the possible conservation of DNA targets recognized by RRBs in *A. thaliana* and *Brassica* species, we performed a multiple protein sequence alignment of DNA-binding GARP-like domain of *A. thaliana* RRBs ARR1, ARR2, ARR10, ARR11, ARR12, ARR13, ARR18, ARR19, ARR20, and ARR21 (Hosoda *et al*., 2002; Lohrmann *et al*., 2001; Mason *et al*., 2005; Sakai *et al*., 2000) and their putative orthologues previously identified in the *Brassica sp.* (Jiang *et al*., 2022; Kaltenegger *et al*., 2018; Liu *et al*., 2014). A high level of conservation was observed with the identity in amino acid sequence ranging from 100% for ARR1, 95.2% for ARR2, and 80.9% for ARR10 to 58.7% in the case of ARR21 (Fig. 6 and Supplementary Fig. S5). Given this high conservation of GARP-like DNA binding domain across the *A. thaliana* and *Brassica* RRBs, it is likely that the *Brassica* RRBs recognize DNA binding motifs similar to those previously described in *A. thaliana* (Hosoda *et al*., 2002; Imamura *et al*., 2003; Sakai *et al*., 2000; Xie *et al*., 2018; Zubo *et al*., 2017).

**Figure 6.**
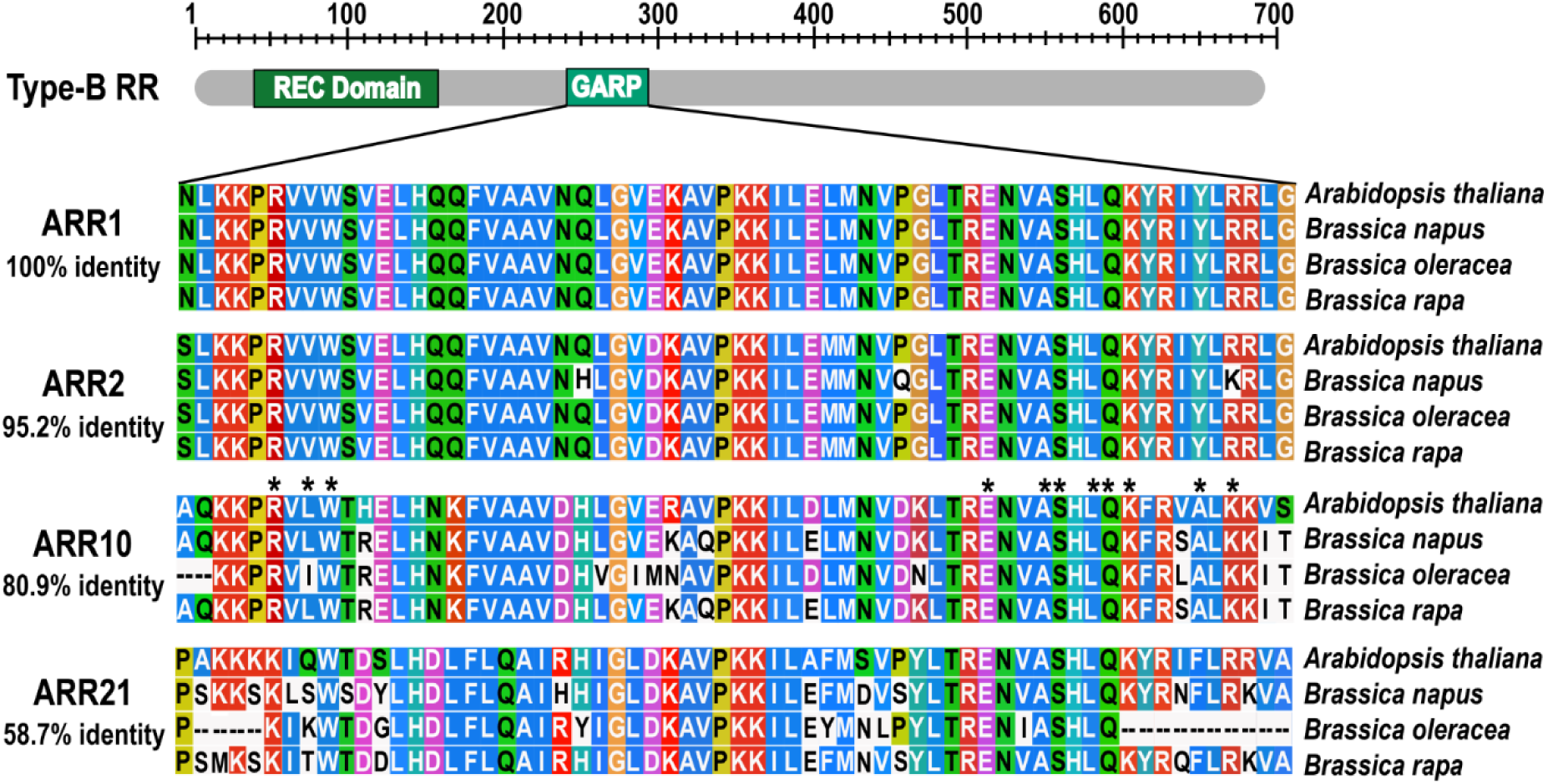
The DNA-binding domain of *A. thaliana* and *Brassica* RRBs show a high level of amino acid conservation. Domain structure of *A. thaliana* and *Brassica* RRBs and alignment of the amino acid sequences of the GARP-like DNA binding domain for the selected RRBs from *A. thaliana* and assayed *Brassica* species. Conserved amino acids are highlighted and the percentage of identity is shown. The CLUSTAL color scheme was used to color the alignment, reflecting the physicochemical properties of amino acids (Kunzmann *et al*., 2020). The asterisk denotes the ARR10 residues that were proposed to directly interact with DNA (Hosoda *et al*., 2002) for a comprehensive list of RRB alignments, refer to Supplementary Figure 5.

To further corroborate this assumption, we utilized the position weight matrices (PWMs) for the *A. thaliana* ARR1 and ARR10 DNA binding sites, retrieved from the ChIP-seq peak sets (Xie *et al*., 2018; Zubo *et al*., 2017) to predict putative RRB binding sites within the *Brassica RRA*s (Fig. 7A). Using this approach, the presence of *Arabidopsis*-like cytokinin-responsive *cis*-elements was predicted in the [−1500; +1] regulatory regions of 62 out of the 66 analyzed *Brassica RRA* genes used in the CK treatment. Similar to *A. thaliana*, these potential *cis*-elements were significantly enriched within the proximal 5’-regulatory regions of *Brassica RRA* genes (within 500 bp upstream of TSS; Fig. 7B). We also observed a moderate correlation between the number of motifs within the [−500;+1] regulatory regions and the magnitude of the transcriptional response to cytokinin, which was statistically significant in *B. napus* and *B. rapa* (Fig. 7C and Supplementary Table S5). This finding further supports the notion of the functional role of *Arabidopsis*-like *cis*-elements in regulating the transcriptional response to cytokinins in the assayed *Brassica* species and suggests the possible role of motif clustering in the response amplification.

**Figure 7.**
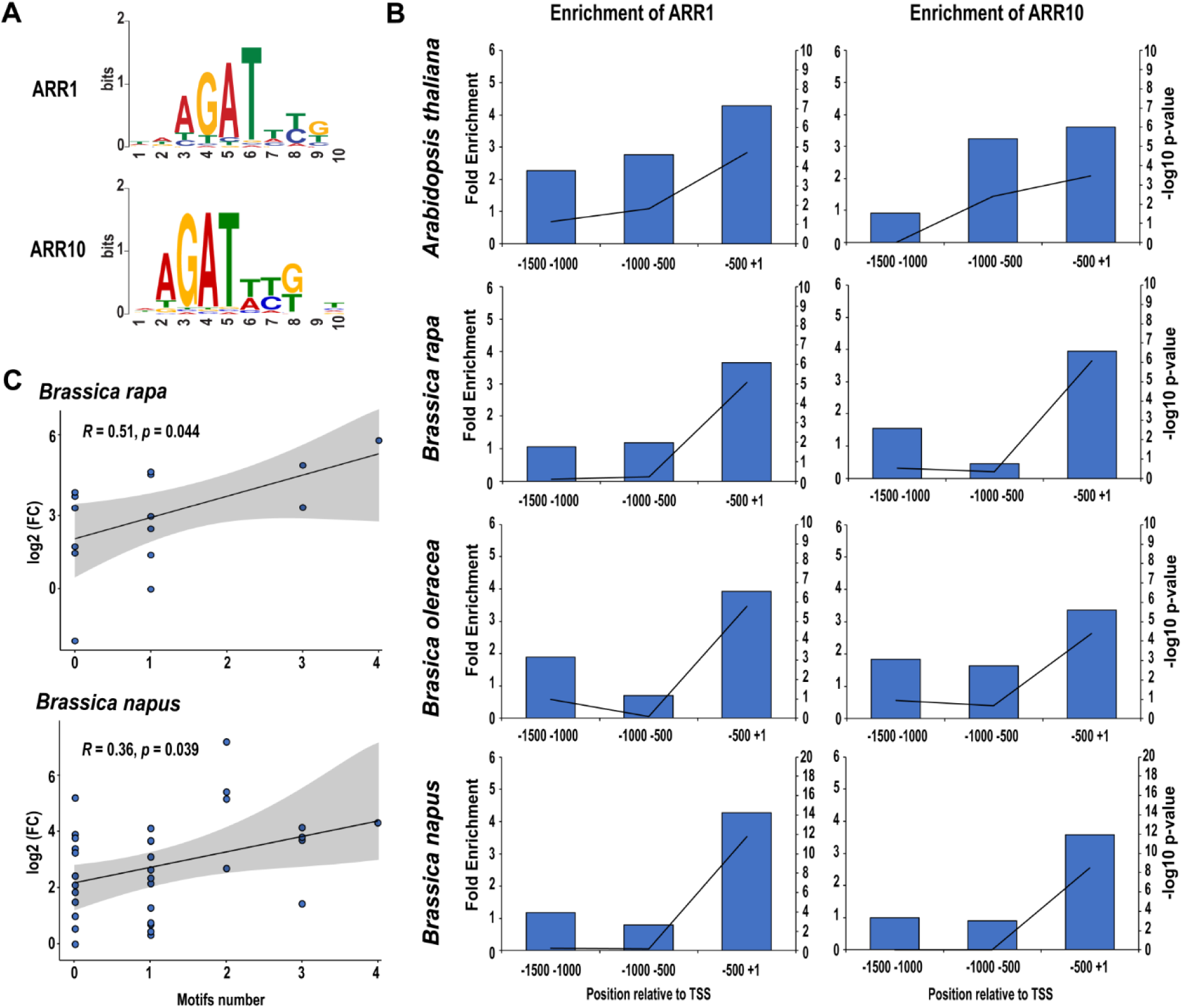
Promoters of Brassica RRAs are enriched for the *Arabidopsis*-like cytokinin-responsive cis-regulatory elements. **(A)** The position weight matrix (PWM) for the ARR1 and ARR10 DNA binding sites in *Arabidopsis thaliana* was retrieved from ChIP-seq peak sets (Xie *et al*., 2018) **(B)** Significant enrichment of ARR1 and ARR10 PWM hits proximally to 5’-regulatory regions of *A. thaliana* and *Brassica RRAs*. Bars represent fold enrichment (left axis) and the line represents log-10 p-value (right axis) **(C)** Significant correlation (Pearson correlation with 95% confidence intervals, shadowed part) between the transcriptional response to cytokinin of *BrRRAs* and *BnRRAs* and the number of cytokinin-responsive motifs present in their promoter regions.

To validate these findings, we utilized a hairy root transformation system (Jedlickova *et al*., 2022) to introduce the cytokinin-responsive reporter (*TCSv2*:3XVENUS) developed in *A. thaliana* by Steiner *et al*. (2020) into *Brassica* species. The *TCSv2* incorporates concatemerized RRB-binding motifs with a distinct arrangement (Fig. 8A) that enhances sensitivity when compared to the original version of the TCS reporter (Zurcher *et al*. (2013). Compared to a mock-treated control, a significant increase in the relative fluorescence intensity was observed after 30 min and 1 hour of the cytokinin treatment in the hairy roots of *B. napus* and *B. rapa*, respectively, carrying the *TCSv2:*3XVENUS (Fig. 8B, C).

**Figure 8.**
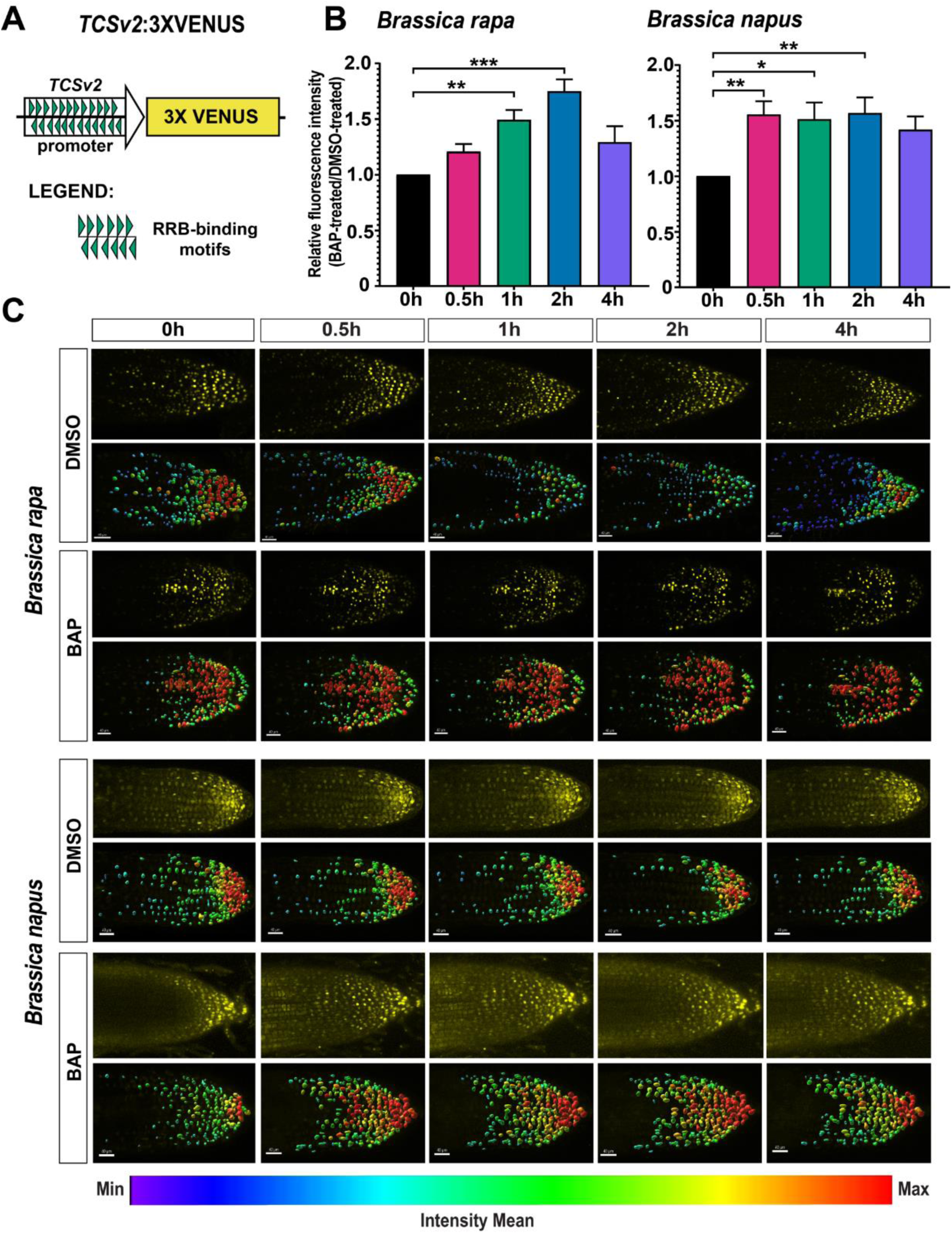
The *Arabidopsis TCSv2*:3XVENUS cytokinin reporter (Steiner *et al*., 2020) is cytokinin-responsive in *B. rapa* and *B. napus*. (A) Scheme of the *TCSv2*:3XVENUS (adapted from (Steiner *et al*., 2020). **(B)** Comparison of the relative fluorescence intensity of *TCSv2*:3XVENUS cytokinin reporter in BAP-treated hairy roots of *B. rapa* and *B. napus* at different time points (0.5, 1, 2, and 4 h) of cytokinin (5 µM BAP) treatment. Means ± standard errors (SE) are shown in the plots. Asterisks indicate statistical significance [p<0.001 (***), p<0.01 (**), and p<0.05 (*), Dunnett’s Test]. **(C)** Representative images of *B. rapa* and *B. napus* hairy root tips treated with DMSO and BAP throughout the treatment period, showing the measured fluorescent signal intensities in a single root (top) and the corresponding image analyzed by IMARIS software (below). Scale bars represent 40 µM.

Taken together, our results strongly suggest that similarly to *Arabidopsis*, the *Brassica* RRBs recognize conserved *cis*-regulatory regions to mediate the cytokinin-induced transcriptional activation of *Brassica RRAs* and possibly other cytokinin-responsive genes within the *Brassica* genomes.

### Cold stress stimulates *RRA* expression in the Brassicaceae family

To assay the possible stress-related regulations of *RRAs* within the Brassicaceae family, the expression profiles of the 66 selected *Brassica RRAs* and the 10 *A. thaliana RRAs* were investigated after exposure to cold, salinity, and osmotic stress. In *A. thaliana*, cold stress rapidly (within 2 h after the stress application) upregulated the expression of several *RRAs*, particularly *ARR6, ARR7,* and *ARR15*. However, the cold-induced upregulation was transient, and the expression of upregulated *RRAs* returned to basal levels after 4 hours of cold exposure. In contrast, we observed gradual repression of *ARR3, ARR8, ARR9, ARR16,* and *ARR17* at 2 h and 4 h of the cold stress application (Fig. 9A and Supplementary Table S6). In *B. rapa,* a greater number of *RRAs* were upregulated in the response to cold, although the induction was delayed when compared to *A. thaliana.* Most *BrRRAs*, except for the non-responsive *BrRRA8a, BrRRA9a, BrRRA9b,* and *BrRRA9c*, exhibited upregulation after 4 h of cold exposure. *BrRRA15a* and *BrRRA15b* showed an earlier response, being upregulated after 2 h of chilling and remaining activated for the 4 hours of the treatment (Fig. 9B and Supplementary Table S6). Also in *B. oleracea*, most of the *BoRRAs* were upregulated by cold stress. Similarly to *A. thaliana*, the response was evident early (2 h) of cold exposure, but compared to the transient upregulation seen in the cold-responsive *A. thaliana RRAs*, the upregulation of *BoRRAs* lasted the entire 4 hours of the treatment. This pattern was observed for *BoRRA6, BoRRA7a, BoRRA7b, BoRRA15a*, and *BoRRA15b* (Fig. 9C and Supplementary Table S6). Also in *B. napus*, we observed prompt upregulation of *RRAs* lasting for the entire 4 hours of the cold treatment. This type of response was apparent for homologs of *ARR6* (*BnARRA6, BnCRRA6*), *ARR7* (*BnARRA7a, BnARRA7b, BnCRRA7a*, and *BnCRRA7b*), and *ARR15* (*BnARRA15a, BnCRRA15b*). Several other *BnRRAs,* including homologs of *ARR3, ARR4, ARR5, ARR8,* and *ARR17,* were also upregulated by cold, but with variable kinetics (Fig. 9D and Supplementary Table S6).

**Figure 9.**
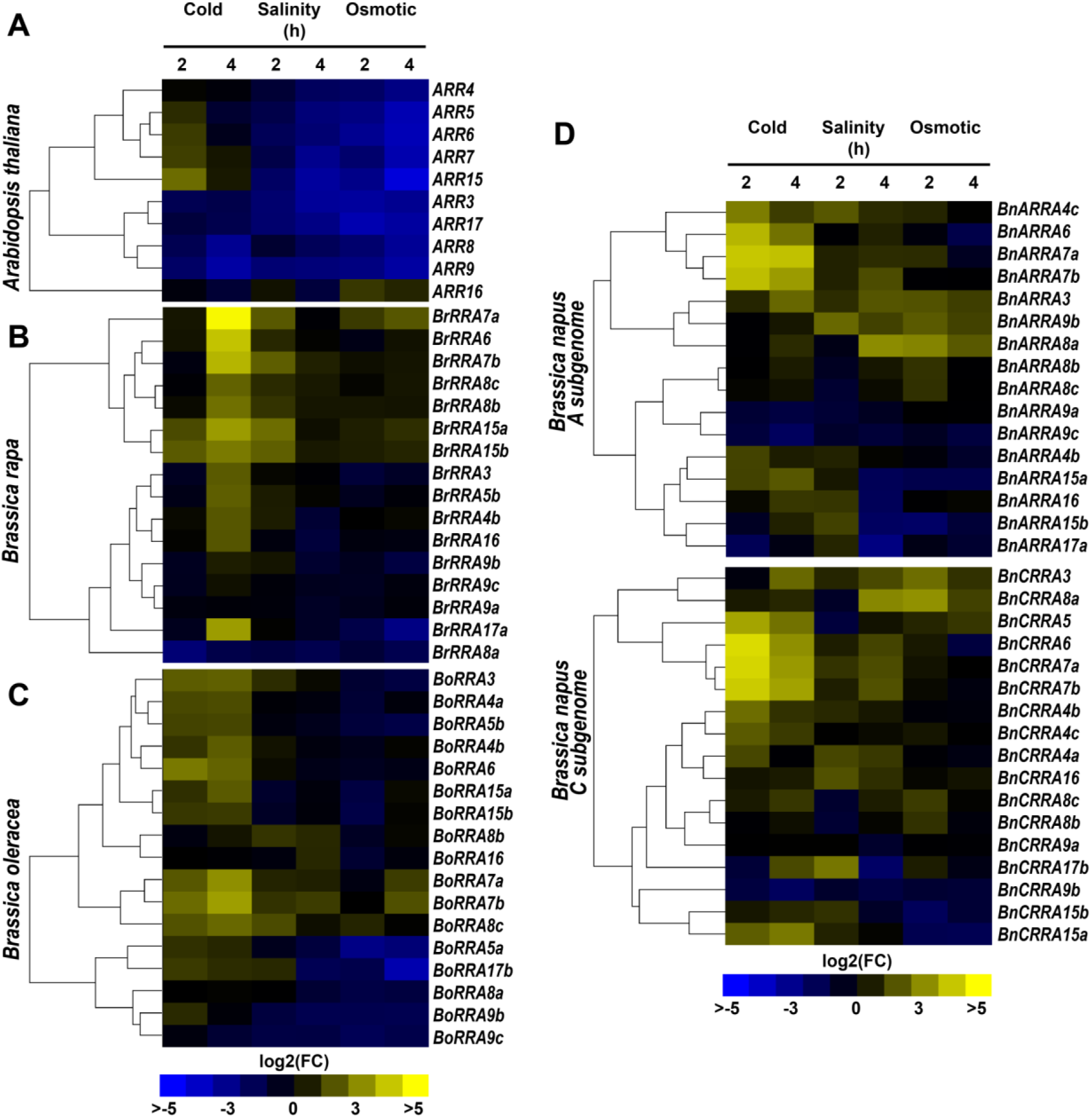
*A. thaliana* and *Brassica RRAs* respond to abiotic stress. Heat maps depicting the expression pattern of RRA genes in the one-week-old seedlings of **(A)** *A. thaliana*, **(B)** *B. rapa*, **(C)** *B. oleracea*, and **(D)** *B. napus* being under cold, salinity, and osmotic stress conditions for 2 and 4 h. The expression data are presented as log2 fold-change normalized to the mock treatment by the delta-delta Ct (Pfaffl, 2004), for the color code see the legend.

To wrap it up, several *RRAs* are upregulated in the response to cold stress in the Brassicaceae family, albeit with slightly different kinetics. Among these, the *ARR6, ARR7, ARR15,* and their *Brassica* homologs appear to represent the core of the common cold-responsive transcriptional signature.

### Salinity and osmotic stress lead to contrasting expression of *A. thaliana* and *Brassica RRAs*

Compared to cytokinin and cold treatment, the majority of *A. thaliana RRAs* exhibited downregulation after exposure to salinity and osmotic stress, except for *ARR16*, which showed upregulation after 2 h of salinity stress (Fig. 9A and Supplementary Table S6). In contrast, several *BrRRAs* were upregulated after 2 h of salinity exposure, particularly the homologs of *ARR6* (*BrRRA6), ARR7* (*BrRRA7a, 7b*), and *ARR15* (*BrRRA15a* and *BrRRA15b*). However, only *BrRRA7b* displayed upregulation when exposed to osmotic stress (Fig. 9B and Supplementary Table S6). In *B. oleracea*, homologs of *ARR7* (*BoRRA7b* and *BoRRA7c)* along with *BoRRA8b* and *BoRRA8c* were upregulated after 2 h of salinity treatment, and this effect persisted up to 4 h except for *BoRRA8b*. In response to osmotic stress, only homologs of *ARR7* (*BoRRA7a* and *BoRRA7b*) were upregulated after 4 h of treatment (Fig. 9C and Supplementary Table S6). In contrast to their diploid ancestors, there were more *RRAs* in *B. napus* that were induced by salinity and/or osmotic stress either after 2 or 4 hours of stress exposure. These included *BnARRA3, BnARRA7a, BnARRA7b*, *BnARRA8a, BnARRA8b, BnARRA8c,* and *BnARRA9b* in the A genome and all *RRAs* from the C-genome except *BnARRA9a* and *BnARRA9b*.

Overall, *RRAs* in Brassicaceae are regulated by salt and osmolarity stresses, displaying various types (up- vs. down-regulation) and dynamics of the response. Compared to *A. thaliana RRAs* being mostly down-regulated, all tested *Brassica* crops exhibited upregulation of *RRAs* not only in the presence of cytokinins but also abiotic stresses. Similar to the cold treatment, homologs of *ARR7* and *ARR15* appear to be a sensitive readout of the response to salinity and high osmolarity in both diploid *Brassicas*, *B. rapa,* and *B. oleracea*. However, particularly in *B. napus,* the response to these stress types seems to be more general, involving a larger number of *RRAs*.

### Cytokinins contribute to the cold stress-induced upregulation of *RRAs* in Brassicaceae

Our gene expression data show the regulation of *RRAs* by abiotic stresses. Utilizing the online database and PlantCARE portal (Lescot *et al*., 2002), several environmental stress-related *cis*-elements were identified in all the promoter sequences of *A. thaliana RRAs,* 16 *BrRRAs* and *BnARRAs*, and 17 *BoRRAs* and *BnCRRAs* (Fig. 10A and Supplementary Table S7). However, the correlation tests between the number of identified stress-related *cis*-elements and the expression of cold-responsive *ARR6*, *ARR7*, *ARR15,* and their *Brassica* homologs after cold exposure did not yield any statistically significant results (Fig. 10B and Supplementary Fig. S6). In an alternative approach, we searched the DAP-seq data (Bartlett *et al*., 2017) to find TFs with potential binding sites in *A. thaliana RRA* promoters. We found 6 such TFs (*AT2G28810, AT3G52440, AT5G56840, ATHB25, ATHB23, ATHB34*); however, the significance of enrichment of their binding sites in the stress-responsive *A. thaliana* and *Brassica RRAs* was low (Supplementary Tables S8-S9). Altogether, our data do not provide any solid evidence supporting the role of the identified stress-related *cis*-regulatory elements in the control of *RRAs* in the Brassicaceae family.

**Figure 10.**
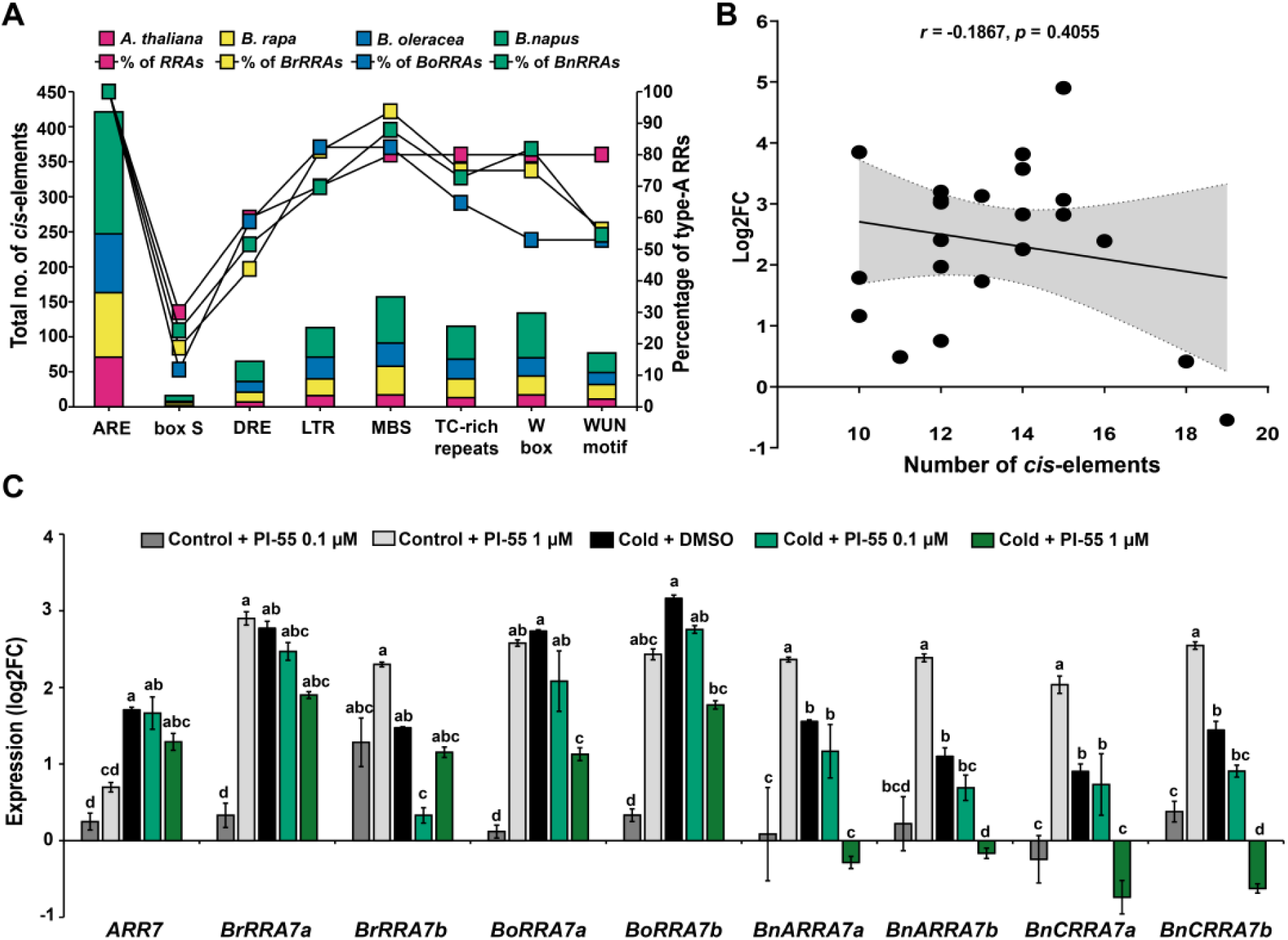
Cytokinins contribute to the cold-induced *ARR7* upregulation. **(A)** Comparison of the number of environmental stress-related *cis*-elements identified in the promoters of *A. thaliana* and *Brassica RRAs* using the PlantCARE program (http://bioinformatics.psb.ugent.be/webtools/plantcare/html/) (Lescot *et al*., 2002) along with the percentage of *RRAs* where these *cis*-elements were found. **(B)** Pearson correlation (with 95% confidence interval, shadowed part) between the transcriptional response of cold-responsive *A. thaliana* and *Brassica RRAs* after 4 hours of cold treatment and the number of environmental *cis*-elements present in their promoter regions. **(C)** Expression of *ARR7* and its homologs after incubation of one-week-old seedlings in media supplemented with either DMSO or cytokinin antagonist PI-55 (0.1 µM/1 µM) and exposure for 4 hours to either cold or control conditions. The expression data are presented as log2 fold-change double normalized by the delta-delta Ct (Pfaffl, 2004) (means ± SE) to the corresponding housekeeping gene (see Materials and Methods) and the control. The different letters indicate variable groups with statistically significant differences (p<0.05, Tukey’s HSD).

To assess the possible involvement of cytokinins in cold stress-mediated upregulation of *RRAs*, we tested the cold response of *ARR7* and its *Brassica* homologs in the presence of anticytokinin PI-55. PI-55 was demonstrated to inhibit the activation of the MSP signaling cascade by competing with cytokinin binding to the CHASE domain of AHKs (Spichal *et al*., 2009). Under control conditions, the treatment with PI-55 led to the induction of all tested *RRAs,* probably due to its previously reported weak cytokinin activity (Spichal *et al*., 2009). However, when applied under low-temperature conditions, PI-55 was able to significantly reduce the cold-induced upregulation of *ARR7* and its *Brassica* homologs. This effect was particularly strong in *B. napus*, where the presence of 1 µM PI-55 completely abolished the upregulation of cold-induced *B. napus ARR7* homologs and led to the drop of gene expression even under the control levels (Fig. 10C).

In conclusion, our findings suggest the existence of a cytokinin-dependent mechanism that contributes to the activation of several *RRAs* in the response to cold stress.

## Discussion

### *Brassica* and *A. thaliana RRAs* reveals close evolutionary relationship

Three rounds of whole genome duplications (WGDs) took place in Brassicaceae after its lineage diverged from monocots but prior to the further divergence within the family (Moghe *et al*., 2014). Kalteneger et al. (2018) proposed the presence of two *RRA* copies (possibly resulting from the ancient ζ or ε WGD event) in the last common ancestor before the divergence of monocots and dicots. Four of the five paralogous *RRA* pairs (ARR6/ARR5, ARR15/ARR7, ARR8/ARR9, and ARR17/ARR16) (Kaltenegger *et al*., 2018) probably originated through the later α WGD event dated to approx. 47 million years ago (Mya). More recently (approx. 25 Mya), an α’ whole-genome triplication (WGT) event took place in the ancestor of *Brassica* species after the divergence from the *Arabidopsis* lineage (Lysak *et al*., 2005; Town *et al*., 2006; Wang *et al*., 2011; Yang *et al*., 2006), leading to the formation of 20 *RRAs* in both *B. oleracea* and *B. rapa*.

An allotetraploid *B. napus* is a result of interspecific hybridization between *B. rapa* and *B. oleracea* (Nagaharu and Nagaharu, 1935; Zhang *et al*., 2016). In accordance with that, the 20 *BnARRAs* identified in the A subgenome and 18 *BnCRRAs* found in the C subgenome, exhibit notable similarity and are mostly syntenic with their counterparts in the *B. rapa* and *B. oleracea* genomes, respectively. Considering the close evolutionary relationships, we used the well-established *A. thaliana RRAs* (*ARRs*) as a reference and numbered the newly identified *B. napus RRAs* according to their (putative) *A. thaliana* orthologues. For the sake of consistency, we extended this type of numbering to the newly identified *BoRRAs* as well as to the previously described *BrRRAs* and *BoRRAs* (Kaltenegger *et al*., 2018). We believe this nomenclature type facilitates comparative analyses within the large gene families of closely related species including the description of gene structure or expression profiles, as we demonstrated in our work. Obviously, different reference species must be used for the monocotyledonous plants that evolved the individual components of (not only) MSP signaling separately (Kaltenegger *et al*., 2018).

### Homologs of *ARR3, ARR6*, and *ARR16* are under evolutionary pressure against multiplication during Brassicaceae evolution

Gene or genome multiplication is an indispensable feature of plant evolution, and gene loss is a frequent fate of newly multiplicated genes (Lynch and Conery, 2000). More specifically, the majority of orthologous groups (approx. 70%) in the common progenitor of recent Brassicaceae species *Raphanus raphanistrum* and *Brassica rapa* experienced losses after the WGT (Moghe *et al*., 2014). Interestingly, genes encoding individual MSP components (i.e., sensor HKs, HPts, and RRs) differ in the extent of gene loss and preservation during evolution. While in the case of HKs, gene loss is a dominant feature, response regulators particularly RRAs are mostly preserved after WGDs (Kaltenegger *et al*., 2018).

In this context, we have rather surprisingly identified three *RRAs*, homologs of *ARR3, ARR6,* and *ARR16,* as singletons in both *B. rapa* and *B. oleracea* (Fig. 3), suggesting evolutionary pressure against the multiplication of those genes. The presence of two copies of the *ARR3, ARR6,* and *ARR16* homologs in *B. napus* (a single copy in each subgenome) might be explained by the recency of the interploidization event. The ability of the gene duplication to be retained seems to be associated with sequence and expression divergence, leading to functional diversification (Moghe *et al*., 2014). In our cytokinin and abiotic stress response assays, we did not observe any strong expression specificity of *ARR3, ARR6,* or *ARR16* and their *Brassica* orthologues, potentially explaining the singleton status of those genes. In *A. thaliana*, some of the *RRAs* were shown to play specific roles in controlling plant growth and development that cannot be solely explained by their functions as redundant cytokinin primary response genes and negative regulators of MSP signaling. To name a few, the ethylene-inducible *ARR3* regulates RAM size (Zdarska *et al*., 2019) and is involved in the cytokinin-independent control over circadian rhythms (Salome *et al*., 2006). ARR6 mediates a negative interaction between abscisic acid and MSP signaling (Huang *et al*., 2017; Wang *et al*., 2011), plays a role in the CLE peptide-mediated inhibition of protoxylem formation (Kondo *et al*., 2011), and regulates pathogen immune response by controlling cell wall composition (Bacete *et al*., 2020). Finally, spatial-specific expression of *ARR16* (and *ARR17)* regulates the hydrotropic bending of the root (Chang *et al*., 2019), and controls stomata formation (Vaten *et al*., 2018) and leaf growth (Efroni *et al*., 2013). Thus, *ARR3, ARR6,* and *ARR16* seem to mediate several key regulatory roles, which might be sensitive to gene dosage. To what extent the *Brassica* homologs of those genes play similar regulatory roles and whether this explains the observed strong negative selection, however, remains to be clarified.

### Cytokinins contribute to abiotic stress-mediated induction of a subset of *RRAs*

The *A. thaliana RRAs* were originally described as cytokinin primary response genes, being rapidly (in order of minutes) induced by exogenous cytokinin treatment (D’Agostino *et al*., 2000). Here, we categorized the *RRAs* based on the kinetics of their cytokinin response into five categories: *(i) fast*, (*ii) fast-transient*, *(iii) no response*, *(iv) later,* and (*v) later transient*. The corresponding response type may reflect certain specificity within MSP signaling (Pekarova *et al*., 2016) with a possible impact on the downstream molecular network underlying the cytokinin cellular responses (Skalak *et al*., 2019). The proportion of individual *RRA* categories varied among tested species with categories *(iv) later* and *(v) later transient* being specific for *Brassica sp*.. However, a subset of *RRAs*, including homologs of *ARR5*, *ARR7,* and *ARR15* [all belonging to the class *(i) fast*] exhibited comparable cytokinin responses in all the tested species. This observation, together with a high level of conservation of the DNA-binding GARP domain of RRBs and the cytokinin responsiveness of *TCSv2* reporter in *B. rapa* and *B. napus*, implies that *RRAs* may share common features and functions within the Brassicaceae family. Interestingly, we observed that a subset of cytokinin-responsive *RRAs* of the category *(i) fast* constitutes a core of the abiotic stress-responsive *RRAs*. While homologs of *ARR6*, *ARR7,* and *ARR15* were cold-responsive, *RRAs* similar to *ARR7* and *ARR15* (together with other *RRAs*, particularly in *B. napus*) seem to be involved also in the response to salinity and high osmolarity in all the tested *Brassicas*, suggesting the existence of common regulatory mechanism. Our finding on the contribution of cytokinin signaling to the cold-mediated regulation of *ARR7* and its *Brassica* homologs is in line with this hypothesis. The (a)biotic stress has been shown to control endogenous hormone levels, including cytokinins, both at the level of biosynthesis and metabolism (Skalak *et al*., 2021) and references therein). This implies that stress-induced upregulation of endogenous cytokinin levels might be a part of cold (and probably other abiotic stress) response in *Brassicaceae*, thus further substantiating the proposed role of plant hormones as a regulatory interface between environmental conditions and intrinsic regulatory pathways controlling individual processes of plant growth and development (Cortleven *et al*., 2019; Ramireddy *et al*., 2014; Skalak *et al*., 2021; Taleski *et al*., 2023; Waadt *et al*., 2022; Yamoune *et al*., 2021).

## Conclusions and future outlines

In summary, our work sheds light on the evolutionary relationships of MSP signaling within the Brassicaceae family. We provide a complete list of the type-A response regulators and their partial molecular characterization in the allotetraploid *B. napus* but also in its parental species, *B. rapa,* and *B. oleracea.* That includes a novel classification reflecting the kinetics of their cytokinin-dependent transcriptional regulation. The conserved occurrence of *ARR3, ARR6,* and *ARR16* as singletons in *A. thaliana*, *B. rapa,* and *B. oleracea* implies the existence of gene-specific negative selection, possibly based on the functional importance and preventing gene multiplication. Several of the *RRAs* exhibited conserved expression patterns in the response to cytokinin and abiotic stresses, implying the presence of common regulatory elements. Our data suggest that cold-mediated induction of *RRAs* demands canonical cytokinin signaling in all tested *Brassica* species, thus emphasizing the importance of cytokinin-regulated MSP in abiotic stress responses. These findings contribute to a nuanced comprehension of the pivotal role of *RRAs* in plant stress responses and open novel avenues for further investigation to uncover the intricate mechanisms guiding plant growth and adaptation, with high potential in applied research.

## Supplementary data

The following supplementary data are available at JXB online.

**Supplementary Table S1.** RRAs from *Arabidopsis thaliana*.

**Supplementary Table S2.** RRAs identified in *Brassica* species.

**Supplementary Table S3.** List of primers used in the study.

**Supplementary Table S4.** Expression profile (log2 fold change) of *RRAs* in *A. thaliana*, *B. rapa*, *B. oleracea*, and *B. napus* after cytokinin treatment.

**Supplementary Table S5.** *Arabidopsis*-like cytokinin-responsive *cis*-elements identified in the promoter regions of *A. thaliana RRAs* and *Brassica RRAs*.

**Supplementary Table S6.** Expression profile (log2 fold change) of *RRAs* in *A. thaliana*, *B. rapa*, *B. oleracea*, and *B. napus* after exposure to abiotic stress.

**Supplementary Table S7.** *In silico* analysis of environmental stress-related *cis*-elements identified using PlantCARE in the promoter regions of the *RRAs* in *A. thaliana*, *B. rapa*, *B. oleracea*, and *B. napus*.

**Supplementary Table S8.** Enrichment of stress-responsive transcription factors in the promoter sequence of *RRAs* in *Arabidopsis thaliana*.

**Supplementary Table S9.** Enrichment of stress-responsive transcription factors in the promoter sequence of *RRAs* in Brassica species.

**Supplementary Figure S1.** Phylogenetic relationship of RRAs in *A. thaliana* and *B. rapa*.

**Supplementary Figure S2.** Phylogenetic relationship of RRAs in *A. thaliana* and *B. oleracea*.

**Supplementary Figure S3.** Phylogenetic relationship of RRAs in *A. thaliana* and those encoded by the A and C subgenomes of *B. napus*.

**Supplementary Figure S4.** The mean expression levels of *RRAs in B. rapa* (*BrRRAs*) and *B. oleracea* (*BoRRAs*) measured from the *Brassica napus* cultivars of the Renewable Industrial Products from Rapeseed (RIPR) diversity panel (Havlickova et al., 2018). The dotted lines indicate the average RPKM values.

**Supplementary Figure S5.** Multiple alignments of GARP-like DNA binding domain of the *A. thaliana* RRBs (ARR11, ARR12, ARR13, ARR14, ARR18, ARR19, and ARR20) *A. thaliana* and their closest homologs from *B. napus, B. oleracea*, and *B. rapa*.

**Supplementary Figure S6.** Pearson correlation (with 95% confidence intervals, shadowed part) between the gene expression of cold-responsive *A. thaliana* and *Brassica* RRAs after 2 hours of cold treatment and the number of environmental stress-related *cis*-elements present in their promoter regions.

## Acknowledgments

We express our gratitude to the Crop Research Institute (Výzkumný ústav rostlinné výroby, v.v.i.) Genebank, Prague, Czech Republic, for providing the *B. napus* (Darmor) seeds. Additionally, we extend our acknowledgment to the core facility CELLIM, which is supported by MEYS CR (LM2023050 Czech-BioImaging), and the Plant Sciences Core Facility of CEITEC Masaryk University for their invaluable technical assistance.

## Author Contributions

JH conceived the research and secured funding, JS and JH coordinated the work. KLNM performed all bioinformatic searches, ranging from BLAST to phylogenetic analysis, RT-qPCR, imaging of the transformed *Brassica* species with CK sensor, statistical analysis, and figure preparation, with assistance from JS. EZ and VD were responsible for promoter analysis, multiple sequence alignment of type B RRs, and figure preparation for this segment. VJ and HSR handled the transformation with a cytokinin sensor and selection of the *Brassica* species. KLNM, JS, EZ, VD, VJ, HSR, LH, IB, and JH wrote and revised the manuscript; all authors read and approved the final manuscript.

## Data availability

All data supporting the findings of this study are available within the paper and its supplementary materials.

## References

Antoniadi I, Novak O, Gelova Z, Johnson A, Plihal O, Simersky R, Mik V, Vain T, Mateo-Bonmati E, Karady M, Pernisova M, Plackova L, Opassathian K, Hejatko J, Robert S, Friml J, Dolezal K, Ljung K, Turnbull C. 2020. Cell-surface receptors enable perception of extracellular cytokinins. Nature communications 11, 4284.

Asakura Y, Hagino T, Ohta Y, Aoki K, Yonekura-Sakakibara K, Deji A, Yamaya T, Sugiyama T, Sakakibara H. 2003. Molecular characterization of His-Asp phosphorelay signaling factors in maize leaves: Implications of the signal divergence by cytokinin-inducible response regulators in the cytosol and the nuclei. Plant Molecular Biology 52, 331–341.

Bacete L, Melida H, Lopez G, Dabos P, Tremousaygue D, Denance N, Miedes E, Bulone V, Goffner D, Molina A. 2020. Arabidopsis Response Regulator 6 (ARR6) Modulates Plant Cell-Wall Composition and Disease Resistance. Molecular plant-microbe interactions : MPMI 33, 767–780.

Bartlett A, O’Malley RC, Huang SSC, Galli M, Nery JR, Gallavotti A, Ecker JR. 2017. Mapping genome-wide transcription-factor binding sites using DAP-seq. Nature Protocols 12, 1659–1672.

Bhaskar A, Paul LK, Sharma E, Jha S, Jain M, Khurana JP. 2021. OsRR6, a type-A response regulator in rice, mediates cytokinin, light and stress responses when over-expressed in Arabidopsis. Plant Physiology and Biochemistry 161, 108–122.

Buechel S, Leibfried A, To JP, Zhao Z, Andersen SU, Kieber JJ, Lohmann JU. 2010. Role of A-type ARABIDOPSIS RESPONSE REGULATORS in meristem maintenance and regeneration. Eur J Cell Biol 89, 279–284.

Cortleven A, Leuendorf JE, Frank M, Pezzetta D, Bolt S, Schmulling T. 2019. Cytokinin action in response to abiotic and biotic stresses in plants. Plant Cell Environ 42, 998–1018.

Cunningham F, Allen JE, Allen J, Alvarez-Jarreta J, Amode M R, Armean Irina M, Austine-Orimoloye O, Azov Andrey G, Barnes I, Bennett R, Berry A, Bhai J, Bignell A, Billis K, Boddu S, Brooks L, Charkhchi M, Cummins C, Da Rin Fioretto L, Davidson C, Dodiya K, Donaldson S, El Houdaigui B, El Naboulsi T, Fatima R, Giron CG, Genez T, Martinez Jose G, Guijarro-Clarke C, Gymer A, Hardy M, Hollis Z, Hourlier T, Hunt T, Juettemann T, Kaikala V, Kay M, Lavidas I, Le T, Lemos D, Marugán JC, Mohanan S, Mushtaq A, Naven M, Ogeh Denye N, Parker A, Parton A, Perry M, Piližota I, Prosovetskaia I, Sakthivel Manoj P, Salam Ahamed Imran A, Schmitt Bianca M, Schuilenburg H, Sheppard D, Pérez-Silva José G, Stark W, Steed E, Sutinen K, Sukumaran R, Sumathipala D, Suner M-M, Szpak M, Thormann A, Tricomi FF, Urbina-Gómez D, Veidenberg A, Walsh Thomas A, Walts B, Willhoft N, Winterbottom A, Wass E, Chakiachvili M, Flint B, Frankish A, Giorgetti S, Haggerty L, Hunt Sarah E, IIsley Garth R, Loveland Jane E, Martin Fergal J, Moore B, Mudge Jonathan M, Muffato M, Perry E, Ruffier M, Tate J, Thybert D, Trevanion Stephen J, Dyer S, Harrison Peter W, Howe Kevin L, Yates Andrew D, Zerbino Daniel R, Flicek P. 2021. Ensembl 2022. Nucleic Acids Research 50, D988–D995.

D’Agostino IB, Deruere J, Kieber JJ. 2000. Characterization of the response of the Arabidopsis response regulator gene family to cytokinin. Plant Physiol 124, 1706–1717.

de Hoon MJL, Imoto S, Nolan J, Miyano S. 2004. Open source clustering software. Bioinformatics 20, 1453–1454.

Duvaud S, Gabella C, Lisacek F, Stockinger H, Ioannidis V, Durinx C. 2021. Expasy, the Swiss Bioinformatics Resource Portal, as designed by its users. Nucleic Acids Research 49, W216–W227.

Edgar RC. 2004. MUSCLE: multiple sequence alignment with high accuracy and high throughput. Nucleic Acids Research 32, 1792–1797.

Efroni I, Han SK, Kim HJ, Wu MF, Steiner E, Birnbaum KD, Hong JC, Eshed Y, Wagner D. 2013. Regulation of Leaf Maturation by Chromatin-Mediated Modulation of Cytokinin Responses. Developmental Cell 24, 438–445.

European Commision. 2019. Oilseeds and protein crops production.

Guénin S, Mauriat M, Pelloux J, Van Wuytswinkel O, Bellini C, Gutierrez L. 2009. Normalization of qRT-PCR data: the necessity of adopting a systematic, experimental conditions-specific, validation of references. Journal of Experimental Botany 60, 487–493.

Havlickova L, He Z, Wang L, Langer S, Harper AL, Kaur H, Broadley MR, Gegas V, Bancroft I. 2018. Validation of an updated Associative Transcriptomics platform for the polyploid crop species Brassica napus by dissection of the genetic architecture of erucic acid and tocopherol isoform variation in seeds. Plant J 93, 181–192.

Heinz S, Benner C, Spann N, Bertolino E, Lin YC, Laslo P, Cheng JX, Murre C, Singh H, Glass CK. 2010. Simple Combinations of Lineage-Determining Transcription Factors Prime cis-Regulatory Elements Required for Macrophage and B Cell Identities. Molecular Cell 38, 576–589.

Hellens RP, Edwards EA, Leyland NR, Bean S, Mullineaux PM. 2000. pGreen: a versatile and flexible binary Ti vector for Agrobacterium-mediated plant transformation. Plant Mol Biol 42, 819–832.

Hendriks KP, Kiefer C, Al-Shehbaz IA, Bailey CD, Huysduynen AHv, Nikolov LA, Nauheimer L, Zuntini AR, German DA, Franzke A, Koch MA, Lysak MA, Toro-Núñez Ó, Özüdoğru B, Invernón VR, Walden N, Maurin O, Hay NM, Shushkov P, Mandáková T, Thulin M, Windham MD, Rešetnik I, Španiel S, Ly E, Pires JC, Harkess A, Neuffer B, Vogt R, Bräuchler C, Rainer H, Janssens SB, Schmull M, Forrest A, Guggisberg A, Zmartzy S, Lepschi BJ, Scarlett N, Stauffer FW, Schönberger I, Heenan P, Baker WJ, Forest F, Mummenhoff K, Lens F. 2022. Global Phylogeny of the Brassicaceae Provides Important Insights into Gene Discordance. bioRxiv, 2022.2009.2001.506188.

Heyl A, Brault M, Frugier F, Kuderova A, Lindner AC, Motyka V, Rashotte AM, Schwartzenberg KV, Vankova R, Schaller GE. 2013. Nomenclature for members of the two-component signaling pathway of plants. Plant Physiology 161, 1063–1065.

Hosoda K, Imamura A, Katoh E, Hatta T, Tachiki M, Yamada H, Mizuno T, Yamazaki T. 2002. Molecular structure of the GARP family of plant Myb-related DNA binding motifs of the Arabidopsis response regulators. Plant Cell 14, 2015–2029.

Howe KL, Achuthan P, Allen J, Allen J, Alvarez-Jarreta J, Amode MR, Armean IM, Azov AG, Bennett R, Bhai J, Billis K, Boddu S, Charkhchi M, Cummins C, Da Rin Fioretto L, Davidson C, Dodiya K, El Houdaigui B, Fatima R, Gall A, Garcia Giron C, Grego T, Guijarro-Clarke C, Haggerty L, Hemrom A, Hourlier T, Izuogu OG, Juettemann T, Kaikala V, Kay M, Lavidas I, Le T, Lemos D, Gonzalez Martinez J, Marugán JC, Maurel T, McMahon AC, Mohanan S, Moore B, Muffato M, Oheh DN, Paraschas D, Parker A, Parton A, Prosovetskaia I, Sakthivel MP, Salam AIA, Schmitt BM, Schuilenburg H, Sheppard D, Steed E, Szpak M, Szuba M, Taylor K, Thormann A, Threadgold G, Walts B, Winterbottom A, Chakiachvili M, Chaubal A, De Silva N, Flint B, Frankish A, Hunt SE, GR II, Langridge N, Loveland JE, Martin FJ, Mudge JM, Morales J, Perry E, Ruffier M, Tate J, Thybert D, Trevanion SJ, Cunningham F, Yates AD, Zerbino DR, Flicek P. 2021. Ensembl 2021.

Hu B, Jin J, Guo AY, Zhang H, Luo J, Gao G. 2015. GSDS 2.0: an upgraded gene feature visualization server.

Huang X, Zhang X, Gong Z, Yang S, Shi Y. 2017. ABI4 represses the expression of type-A ARRs to inhibit seed germination in Arabidopsis. Plant J 89, 354–365.

Hwang I, Chen HC, Sheen J. 2002. Two-component signal transduction pathways in Arabidopsis. Plant Physiol 129, 500–515.

Hwang I, Sheen J. 2001. Two-component circuitry in Arabidopsis cytokinin signal transduction. Nature 413, 383–389.

Chang J, Li X, Fu W, Wang J, Yong Y, Shi H, Ding Z, Kui H, Gou X, He K, Li J. 2019. Asymmetric distribution of cytokinins determines root hydrotropism in Arabidopsis thaliana. Cell Res 29, 984–993.

Chao JT, Kong YZ, Wang Q, Sun YH, Gong DP, Lv J, Liu GS. 2015. MapGene2Chrom, a tool to draw gene physical map based on Perl and SVG languages.

Chen C, Chen H, Zhang Y, Thomas HR, Frank MH, He Y, Xia R. 2020. TBtools: An Integrative Toolkit Developed for Interactive Analyses of Big Biological Data. Molecular Plant 13, 1194–1202.

Chen H, Wang T, He X, Cai X, Lin R, Liang J, Wu J, King G, Wang X. 2021. BRAD V3.0: an upgraded Brassicaceae database. Nucleic Acids Research 50, D1432–D1441.

Cheng F, Wu J, Fang L, Wang X. 2012. Syntenic gene analysis between Brassica rapa and other Brassicaceae species. Frontiers in plant science 3.

Cheng F, Wu J, Wang X. 2014. Genome triplication drove the diversification of Brassica plants. Horticulture Research 1, 14024.

Imamura A, Hanaki N, Umeda H, Nakamura A, Suzuki T, Ueguchi C, Mizuno T. 1998. Response regulators implicated in His-to-Asp phosphotransfer signaling in Arabidopsis. Proceedings of the National Academy of Sciences of the United States of America 95, 2691–2696.

Imamura A, Kiba T, Tajima Y, Yamashino T, Mizuno T. 2003. In vivo and in vitro characterization of the ARR11 response regulator implicated in the His-to-Asp phosphorelay signal transduction in Arabidopsis thaliana. Plant and Cell Physiology 44, 122–131.

Inoue T, Higuchi M, Hashimoto Y, Seki M, Kobayashi M, Kato T, Tabata S, Shinozaki K, Kakimoto T. 2001. Identification of CRE1 as a cytokinin receptor from Arabidopsis. Nature 409, 1060–1063.

Ito Y, Kurata N. 2006. Identification and characterization of cytokinin-signalling gene families in rice. Gene 382, 57–65.

Jain M, Tyagi AK, Khurana JP. 2006. Molecular characterization and differential expression of cytokinin-responsive type-A response regulators in rice (Oryza sativa). BMC Plant Biology 6, 1.

Jedlickova V, Macova K, Stefkova M, Butula J, Stavenikova J, Sedlacek M, Robert HS. 2022. Hairy root transformation system as a tool for CRISPR/Cas9-directed genome editing in oilseed rape (Brassica napus). Frontiers in plant science 13, 919290.

Jeon J, Kim NY, Kim S, Kang NY, Novak O, Ku SJ, Cho C, Lee DJ, Lee EJ, Strnad M, Kim J. 2010. A subset of cytokinin two-component signaling system plays a role in cold temperature stress response in Arabidopsis. J Biol Chem 285, 23371–23386.

Jiang JJ, Li N, Chen WJ, Wang Y, Rong H, Xie T, Wang YP. 2022. Genome-Wide Analysis of the Type-B Authentic Response Regulator Gene Family in Brassica napus. Genes (Basel) 13.

Kaltenegger E, Leng S, Heyl A. 2018. The effects of repeated whole genome duplication events on the evolution of cytokinin signaling pathway. BMC Evol Biol 18, 76.

Kang NY, Cho C, Kim NY, Kim J. 2012. Cytokinin receptor-dependent and receptor-independent pathways in the dehydration response of Arabidopsis thaliana. Journal of Plant Physiology 169, 1382–1391.

Karan R, Singla-Pareek SL, Pareek A. 2009. Histidine kinase and response regulator genes as they relate to salinity tolerance in rice. Functional & integrative genomics 9, 411–417.

Kieber JJ, Schaller GE. 2018. Cytokinin signaling in plant development. Development 145.

Kondo Y, Hirakawa Y, Kieber JJ, Fukuda H. 2011. CLE peptides can negatively regulate protoxylem vessel formation via cytokinin signaling. Plant Cell Physiol 52, 37–48.

Kubiasova K, Montesinos JC, Samajova O, Nisler J, Mik V, Semeradova H, Plihalova L, Novak O, Marhavy P, Cavallari N, Zalabak D, Berka K, Dolezal K, Galuszka P, Samaj J, Strnad M, Benkova E, Plihal O, Spichal L. 2020. Cytokinin fluoroprobe reveals multiple sites of cytokinin perception at plasma membrane and endoplasmic reticulum. Nature communications 11, 4285.

Kuderova A, Gallova L, Kuricova K, Nejedla E, Curdova A, Micenkova L, Plihal O, Smajs D, Spichal L, Hejatko J. 2015. Identification of AHK2- and AHK3-like cytokinin receptors in Brassica napus reveals two subfamilies of AHK2 orthologues. Journal of Experimental Botany 66, 339–353.

Kumar A, Sharma P, Thomas L, Agnihotri A, Banga S. 2009. Canola cultivation in India: scenario and future strategy. 16th Australian research assembly on Brassicas. Ballarat, Victoria, 0–5.

Kumar S, Stecher G, Tamura K. 2016. MEGA7: Molecular Evolutionary Genetics Analysis Version 7.0 for Bigger Datasets. Molecular Biology and Evolution 33, 1870–1874.

Kunzmann P, Mayer BE, Hamacher K. 2020. Substitution matrix based color schemes for sequence alignment visualization. BMC Bioinformatics 21, 209.

Kyoto University Bioinformatics Center. 2015. MOTIF: Searching Protein Sequence Motifs.

Lee DJ, Kim S, Ha YM, Kim J. 2008. Phosphorylation of Arabidopsis response regulator 7 (ARR7) at the putative phospho-accepting site is required for ARR7 to act as a negative regulator of cytokinin signaling. Planta 227, 577–587.

Leibfried A, To JP, Busch W, Stehling S, Kehle A, Demar M, Kieber JJ, Lohmann JU. 2005. WUSCHEL controls meristem function by direct regulation of cytokinin-inducible response regulators. Nature 438, 1172–1175.

Lescot M, Déhais P, Thijs G, Marchal K, Moreau Y, Van de Peer Y, Rouzé P, Rombauts S. 2002. PlantCARE, a database of plant cis-acting regulatory elements and a portal to tools for in silico analysis of promoter sequences. Nucleic Acids Research 30, 325–327.

Liu ZN, Zhang M, Kong LJ, Lv YX, Zou MH, Lu G, Cao JS, Yu XL. 2014. Genome-Wide Identification, Phylogeny, Duplication, and Expression Analyses of Two-Component System Genes in Chinese Cabbage (Brassica rapa ssp pekinensis). DNA Research 21, 379–396.

Lohrmann J, Sweere U, Zabaleta E, Baurle I, Keitel C, Kozma-Bognar L, Brennicke A, Schafer E, Kudla J, Harter K. 2001. The response regulator ARR2: a pollen-specific transcription factor involved in the expression of nuclear genes for components of mitochondrial Complex I in Arabidopsis. Molecular Genetics and Genomics 265, 2–13.

Lynch M, Conery JS. 2000. The evolutionary fate and consequences of duplicate genes. Science 290, 1151–1155.

Lysak MA, Koch MA, Pecinka A, Schubert I. 2005. Chromosome triplication found across the tribe Brassiceae. Genome Research 15, 516–525.

Madeira FA-O, Pearce MA-O, Tivey ARN, Basutkar P, Lee J, Edbali O, Madhusoodanan N, Kolesnikov A, Lopez RA-O. 2022. Search and sequence analysis tools services from EMBL-EBI in 2022.

Mason MG, Mathews DE, Argyros DA, Maxwell BB, Kieber JJ, Alonso JM, Ecker JR, Schaller GE. 2005. Multiple type-B response regulators mediate cytokinin signal transduction in Arabidopsis. Plant Cell 17, 3007–3018.

Moghe GD, Hufnagel DE, Tang H, Xiao Y, Dworkin I, Town CD, Conner JK, Shiu SH. 2014. Consequences of Whole-Genome Triplication as Revealed by Comparative Genomic Analyses of the Wild Radish Raphanus raphanistrum and Three Other Brassicaceae Species. Plant Cell 26, 1925–1937.

Mochida K, Yoshida T, Sakurai T, Yamaguchi-Shinozaki K, Shinozaki K, Tran LS. 2010. Genome-wide analysis of two-component systems and prediction of stress-responsive two-component system members in soybean. DNA research : an international journal for rapid publication of reports on genes and genomes 17, 303–324.

Morinaga T. 1929. lnterspecific Hybridization in Brassica I. The Cytology of Fl Hybrids of B. Napella and Various Other Species with 10 Chromosomes. Cytologia 1, 16–27.

Muller B, Sheen J. 2007. Advances in cytokinin signaling. Science 318, 68–69.

Muller B, Sheen J. 2008. Cytokinin and auxin interaction in root stem-cell specification during early embryogenesis. Nature 453, 1094–1097.

Nagaharu U, Nagaharu N. 1935. Genome analysis in Brassica with special reference to the experimental formation of B. napus and peculiar mode of fertilization. Jpn. J. Bot. 7, 389–452.

Nikolov LA, Shushkov P, Nevado B, Gan X, Al-Shehbaz IA, Filatov D, Bailey CD, Tsiantis M. 2019. Resolving the backbone of the Brassicaceae phylogeny for investigating trait diversity. New Phytol 222, 1638–1651.

Okonechnikov K, Golosova O, Fursov M, team tU. 2012a. Unipro UGENE: a unified bioinformatics toolkit. Bioinformatics 28, 1166–1167.

Okonechnikov K, Golosova O, Fursov M, Team U. 2012b. Unipro UGENE: a unified bioinformatics toolkit. Bioinformatics 28, 1166–1167.

Pareek A, Singh A, Kumar M, Kushwaha HR, Lynn AM, Singla-Pareek SL. 2006. Whole-genome analysis of Oryza sativa reveals similar architecture of two-component signaling machinery with Arabidopsis. Plant Physiology 142, 380–397.

Pekarova B, Szmitkowska A, Dopitova R, Degtjarik O, Zidek L, Hejatko J. 2016. Structural Aspects of Multistep Phosphorelay-Mediated Signaling in Plants. Mol Plant 9, 71–85.

Pfaffl MW. 2004. Quantification strategies in real-time PCR. In: Bustin SA, ed. A-Z of quantitative PCR. La Jolla, CA, USA.

Quinlan AR, Hall IM. 2010. BEDTools: a flexible suite of utilities for comparing genomic features. Bioinformatics 26, 841–842.

Ramireddy E, Chang L, Schmulling T. 2014. Cytokinin as a mediator for regulating root system architecture in response to environmental cues. Plant Signal Behav 9, e27771.

Rathore SS, Babu S, Shekhawat K, Singh VK, Upadhyay PK, Singh RK, Raj R, Singh H, Zaki FM. 2022. Oilseed Brassica Species Diversification and Crop Geometry Influence the Productivity, Economics, and Environmental Footprints under Semi-Arid Regions. Sustainability 14, 2230.

Saitou N, Nei M. 1987. The neighbor-joining method: a new method for reconstructing phylogenetic trees. Molecular Biology and Evolution 4, 406–425.

Sakai H, Aoyama T, Oka A. 2000. Arabidopsis ARR1 and ARR2 response regulators operate as transcriptional activators. The Plant journal : for cell and molecular biology 24, 703–711.

Saldanha AJ. 2004. Java Treeview—extensible visualization of microarray data. Bioinformatics 20, 3246–3248.

Salome PA, To JP, Kieber JJ, McClung CR. 2006. Arabidopsis response regulators ARR3 and ARR4 play cytokinin-independent roles in the control of circadian period. Plant Cell 18, 55–69.

Sharan A, Soni P, Nongpiur RC, Singla-Pareek SL, Pareek A. 2017. Mapping the ‘Two-component system’ network in rice. Scientific reports 7, 9287.

Shi Y, Tian S, Hou L, Huang X, Zhang X, Guo H, Yang S. 2012. Ethylene signaling negatively regulates freezing tolerance by repressing expression of CBF and type-A ARR genes in Arabidopsis. Plant Cell 24, 2578–2595.

Schaller GE, Kieber JJ, Shiu SH. 2008. Two-component signaling elements and histidyl-aspartyl phosphorelays. Arabidopsis Book 6, e0112.

Schneider CA, Rasband WS, Eliceiri KW. 2012. NIH Image to ImageJ: 25 years of image analysis. Nature Methods 9, 671–675.

Skalak J, Nicolas KL, Vankova R, Hejatko J. 2021. Signal Integration in Plant Abiotic Stress Responses via Multistep Phosphorelay Signaling. Frontiers in plant science 12, 644823.

Skalak J, Vercruyssen L, Claeys H, Hradilova J, Cerny M, Novak O, Plackova L, Saiz-Fernandez I, Skalakova P, Coppens F, Dhondt S, Koukalova S, Zouhar J, Inze D, Brzobohaty B. 2019. Multifaceted activity of cytokinin in leaf development shapes its size and structure in Arabidopsis. Plant J 97, 805–824.

Spichal L, Werner T, Popa I, Riefler M, Schmulling T, Strnad M. 2009. The purine derivative PI-55 blocks cytokinin action via receptor inhibition. FEBS J 276, 244–253.

Steiner E, Israeli A, Gupta R, Shwartz I, Nir I, Leibman-Markus M, Tal L, Farber M, Amsalem Z, Ori N, Muller B, Bar M. 2020. Characterization of the cytokinin sensor TCSv2 in arabidopsis and tomato. Plant Methods 16, 152.

Sun L, Lv L, Zhao J, Hu M, Zhang Y, Zhao Y, Tang X, Wang P, Li Q, Chen X, Li H, Zhang Y. 2022. Genome-wide identification and expression analysis of the TaRRA gene family in wheat (Triticum aestivum L.). Frontiers in plant science 13, 1006409.

Taleski M, Chapman K, Novak O, Schmulling T, Frank M, Djordjevic MA. 2023. CEP peptide and cytokinin pathways converge on CEPD glutaredoxins to inhibit root growth. Nature communications 14, 1683.

Taniguchi M, Kiba T, Sakakibara H, Ueguchi C, Mizuno T, Sugiyama T. 1998. Expression of Arabidopsis response regulator homologs is induced by cytokinins and nitrate. FEBS Letters 429, 259–262.

To JP, Haberer G, Ferreira FJ, Deruere J, Mason MG, Schaller GE, Alonso JM, Ecker JR, Kieber JJ. 2004. Type-A Arabidopsis response regulators are partially redundant negative regulators of cytokinin signaling. Plant Cell 16, 658–671.

Touzet H, Varré J-S. 2007. Efficient and accurate P-value computation for Position Weight Matrices. Algorithms for Molecular Biology 2, 15.

Town CD, Cheung F, Maiti R, Crabtree J, Haas BJ, Wortman JR, Hine EE, Althoff R, Arbogast TS, Tallon LJ, Vigouroux M, Trick M, Bancroft I. 2006. Comparative genomics of Brassica oleracea and Arabidopsis thaliana reveal gene loss, fragmentation, and dispersal after polyploidy. Plant Cell 18, 1348–1359.

Tran LS, Urao T, Qin F, Maruyama K, Kakimoto T, Shinozaki K, Yamaguchi-Shinozaki K. 2007. Functional analysis of AHK1/ATHK1 and cytokinin receptor histidine kinases in response to abscisic acid, drought, and salt stress in Arabidopsis. Proc Natl Acad Sci U S A 104, 20623–20628.

Tsai YC, Weir NR, Hill K, Zhang W, Kim HJ, Shiu SH, Schaller GE, Kieber JJ. 2012. Characterization of genes involved in cytokinin signaling and metabolism from rice. Plant Physiology 158, 1666–1684.

Urao T, Yakubov B, Yamaguchi-Shinozaki K, Shinozaki K. 1998. Stress-responsive expression of genes for two-component response regulator-like proteins in Arabidopsis thaliana. FEBS Lett 427, 175–178.

Vaten A, Soyars CL, Tarr PT, Nimchuk ZL, Bergmann DC. 2018. Modulation of Asymmetric Division Diversity through Cytokinin and SPEECHLESS Regulatory Interactions in the Arabidopsis Stomatal Lineage. Dev Cell 47, 53–66 e55.

Waadt R, Seller CA, Hsu PK, Takahashi Y, Munemasa S, Schroeder JI. 2022. Plant hormone regulation of abiotic stress responses. Nat Rev Mol Cell Biol 23, 680–694.

Wang WC, Lin TC, Kieber J, Tsai YC. 2019. Response Regulator 9 and 10 Negatively Regulate Salinity Tolerance in Rice. Plant Cell Physiol.

Wang Y, Li L, Ye T, Zhao S, Liu Z, Feng YQ, Wu Y. 2011. Cytokinin antagonizes ABA suppression to seed germination of Arabidopsis by downregulating ABI5 expression. Plant J 68, 249–261.

Werner T, Schmulling T. 2009. Cytokinin action in plant development. Curr Opin Plant Biol 12, 527–538.

Xie M, Chen H, Huang L, O’Neil RC, Shokhirev MN, Ecker JR. 2018. A B-ARR-mediated cytokinin transcriptional network directs hormone cross-regulation and shoot development. Nature communications 9, 1604.

Yamoune A, Cuyacot AR, Zdarska M, Hejatko J. 2021. Hormonal orchestration of root apical meristem formation and maintenance in Arabidopsis. J Exp Bot 72, 6768–6788.

Yang TJ, Kim JS, Kwon SJ, Lim KB, Choi BS, Kim JA, Jin M, Park JY, Lim MH, Kim HI, Lim YP, Kang JJ, Hong JH, Kim CB, Bhak J, Bancroft I, Parka BS. 2006. Sequence-level analysis of the diploidization process in the triplicated FLOWERING LOCUS C region of Brassica rapa. Plant Cell 18, 1339–1347.

Yates AA-O, Allen J, Amode RM, Azov AG, Barba M, Becerra A, Bhai J, Campbell LI, Carbajo Martinez M, Chakiachvili M, Chougule K, Christensen M, Contreras-Moreira B, Cuzick AA-O, Da Rin Fioretto L, Davis PA-O, De Silva NA-O, Diamantakis S, Dyer S, Elser J, Filippi CV, Gall A, Grigoriadis D, Guijarro-Clarke C, Gupta P, Hammond-Kosack KA-OX, Howe KA-O, Jaiswal PA-O, Kaikala V, Kumar V, Kumari S, Langridge N, Le T, Luypaert M, Maslen GL, Maurel T, Moore B, Muffato M, Mushtaq A, Naamati G, Naithani SA-O, Olson A, Parker A, Paulini M, Pedro H, Perry EA-O, Preece J, Quinton-Tulloch M, Rodgers F, Rosello M, Ruffier MA-O, Seager JA-OX, Sitnik V, Szpak M, Tate J, Tello-Ruiz MK, Trevanion SJ, Urban MA-O, Ware D, Wei S, Williams G, Winterbottom A, Zarowiecki M, Finn RA-O, Flicek PA-O. 2022. Ensembl Genomes 2022: an expanding genome resource for non-vertebrates.

Zdarska M, Cuyacot AR, Tarr PT, Yamoune A, Szmitkowska A, Hrdinova V, Gelova Z, Meyerowitz EM, Hejatko J. 2019. ETR1 Integrates Response to Ethylene and Cytokinins into a Single Multistep Phosphorelay Pathway to Control Root Growth. Mol Plant 12, 1338–1352.

Zhang X, Liu T, Li X, Duan M, Wang J, Qiu Y, Wang H, Song J, Shen D. 2016. Interspecific hybridization, polyploidization and backcross of Brassica oleracea var. alboglabra with B. rapa var. purpurea morphologically recapitulate the evolution of Brassica vegetables. Scientific reports 6, 18618.

Zhao Z, Andersen SU, Ljung K, Dolezal K, Miotk A, Schultheiss SJ, Lohmann JU. 2010. Hormonal control of the shoot stem-cell niche. Nature 465, 1089–1092.

Zubo YO, Blakley IC, Yamburenko MV, Worthen JM, Street IH, Franco-Zorrilla JM, Zhang W, Hill K, Raines T, Solano R, Kieber JJ, Loraine AE, Schaller GE. 2017. Cytokinin induces genome-wide binding of the type-B response regulator ARR10 to regulate growth and development in Arabidopsis. Proc Natl Acad Sci U S A 114, E5995–E6004.

Zuckerkandl E, Pauling L. 1965. Evolutionary Divergence and Convergence in Proteins. In: Bryson V, Vogel HJ, eds. *Evolving Genes and Proteins*: Academic Press, 97–166.

Zurcher E, Muller B. 2016. Cytokinin Synthesis, Signaling, and Function--Advances and New Insights. Int Rev Cell Mol Biol 324, 1–38.

Zurcher E, Tavor-Deslex D, Lituiev D, Enkerli K, Tarr PT, Muller B. 2013. A robust and sensitive synthetic sensor to monitor the transcriptional output of the cytokinin signaling network in planta. Plant Physiology 161, 1066–1075.

